# The coordinate actions of calcineurin and Hog1 mediate the response to cellular stress through multiple nodes of the cell cycle network

**DOI:** 10.1101/612416

**Authors:** Cassandra M. Leech, Mackenzie J. Flynn, Heather E. Arsenault, Jianhong Ou, Haibo Liu, Lihua Julie Zhu, Jennifer A. Benanti

**Affiliations:** Department of Molecular, Cell and Cancer Biology, University of Massachusetts Medical School, Worcester, MA 01605; Program in Bioinformatics and Integrative Biology, Program in Molecular Medicine, University of Massachusetts Medical School, Worcester, MA 01605

## Abstract

Upon exposure to environmental stressors, cells transiently arrest the cell cycle while they adapt and restore homeostasis. A challenge for all cells is to distinguish between diverse stress signals and coordinate the appropriate adaptive response with cell cycle arrest. Here we investigate the role of the stress-activated phosphatase calcineurin (CN) in this process and show that CN utilizes multiple pathways to control the cell cycle. Upon activation, CN inhibits transcription factors (TFs) that regulate the G1/S transition through activation of the stress-activated MAPK Hog1. In contrast, CN inactivates G2/M TFs through a combination of Hog1-dependent and -independent mechanisms. These findings demonstrate that CN and Hog1 act in a coordinated manner at multiple nodes of the cell cycle-regulatory network to rewire gene expression and arrest cells in response to stress. Our results suggest that crosstalk between CN and stress-activated MAPKs helps cells tailor their adaptive responses to specific stressors.

## Introduction

Cells must constantly monitor their environment and correctly interpret extracellular signals such that they grow and divide only in favorable conditions. When cells attempt to divide in unfavorable conditions this often results in cell death. Therefore, the signaling pathways that sense and interpret changes in the environment are critical. Cells must detect and distinguish among a wide array of environmental stressors including oxidative stress, temperature, DNA damage, and changes in pH or osmolarity. In each of these cases, cells transiently arrest the cell cycle, while also promoting stress-specific changes in post-translational modifications and gene expression that allow cells to adapt to their new environment (Ho and Gasch, 2015). To mount specific responses against all of these diverse insults, cells utilize multiple signaling pathways that respond to different inputs. However, the mechanisms by which different stress-response pathways work together to coordinate cell cycle arrest and adaptation to specific stressors is not well understood.

The cell cycle is driven by a transcriptional program that is orchestrated by an interconnected network of transcription factors (TFs). Cyclical transcription established by these TFs ensures that cell cycle regulators are expressed precisely at the times their functions are needed and promotes unidirectional progression through the cell cycle (Bertoli et al., 2013; Haase and Wittenberg, 2014; Sadasivam and DeCaprio, 2013). Many environmental stressors trigger checkpoints that inhibit the activities of these TFs and impair cell cycle progression. This occurs at both the G1/S restriction point in mammals (START in yeast), when cells decide whether or not to commit to DNA replication, and at the G2/M transition, before cells proceed into mitosis (Morgan, 2007).

A critical mediator of the cellular stress response is the calcium/calmodulin-activated phosphatase calcineurin (CN), which is essential for cells to survive numerous environmental insults. In mammals, oxidative stress and nutrient starvation promote the release of lysosomal Ca^2+^, which activates CN, leading to increased lysosome biogenesis and autophagy (Medina et al., 2015; Zhang et al., 2016). Similarly, a number of environmental stressors, including alkaline pH, toxic ions, and cell wall stress, trigger an influx of cytoplasmic Ca^2+^ that activates CN in budding yeast (Cyert and Philpott, 2013). The best studied downstream effector of CN in this system is the TF Crz1, which upon dephosphorylation by CN translocates to the nucleus and activates transcription of approximately 150 target genes (Yoshimoto et al., 2002). Crz1 targets include many regulators that feedback and shut off Ca^2+^-signaling and the CN response, which promotes adaptation to the stress. In addition to activating gene expression via Crz1, CN regulates a number of processes that change the physiology of the cell to allow it to cope with stress including protein trafficking, transcription, polarized growth, and others (Cyert and Philpott, 2013; Goldman et al., 2014).

A few cell cycle regulators have also been identified as CN targets, suggesting that CN may help coordinate the stress response with cell cycle arrest (Goldman et al., 2014; Mizunuma et al., 2001). Inactivation of the S-phase specific transcriptional activator Hcm1 by CN leads to decreased expression of its target genes, which include TFs that act at a later stage in the cell cycle (Arsenault et al., 2015). However, it is not known if CN impairs cell cycle-regulated gene expression solely through inactivation of Hcm1, or if CN regulates the TF network through additional mechanisms.

Here, we sought to obtain a network level view of how CN impacts the cell cycle to better understand the mechanisms by which cells respond to cellular stresses. To this end, we analyzed the temporal response of the cell cycle-regulated transcriptome in response to CN activation and characterized the pathways mediating the CN response at each node of the cell cycle. Remarkably, we find that CN downregulates targets of multiple cell cycle TFs through distinct mechanisms. We find that CN blocks expression of G1/S genes by stimulating the activity of the stress-activated MAPK Hog1, an established inhibitor of G1/S TFs (Bellí et al., 2001; González-Novo et al., 2015). In contrast, CN inactivates G2/M TFs through both Hog1-dependent and -independent mechanisms. In this way, cells tailor their response to Ca^2+^-stress by coordinating the activation of CN and Hog1. Together, CN and Hog1 trigger widespread rewiring of the cell cycle-regulated transcriptional program as well as a transient cell cycle arrest that enables cells to rapidly respond to and recover from environmental stress.

## Results

### CN downregulates multiple clusters of cell cycle genes

To understand the dynamics of cell cycle regulation by CN as cells respond and adapt to stress, we followed the cell cycle distribution of a population of asynchronous cells over time following CN activation. Cells were treated with CaCl_2_ to elicit strong activation of CN, following pre-treatment with either the CN inhibitor FK506 or buffer alone to identify CN-specific changes. In addition, to lengthen the window of time before cells adapted to the CaCl_2_ stress, experiments were performed in cells lacking the CN target Crz1. In these conditions, the addition of CaCl_2_ arrested cells by 30 minutes, which was evident by the depletion of cells in S-phase (Figures 1A and S1A). Interestingly, this arrest occurred in both control (ET buffer) and FK506-treated cells, indicating that the initiation of cell cycle arrest is CN-independent. However, at later time points, FK506-treated cells resumed cycling while control cells remained arrested (Figure 1A). A similar pattern of cell cycle arrest also occurred in wild-type (*CRZ1* proficient) cells, although cells resumed cycling more quickly, as expected (Figure S1B). These data demonstrate that the transient cell cycle arrest induced by CaCl_2_ occurs independently of CN, however CN is required to maintain the arrest over time.

**Figure 1.**
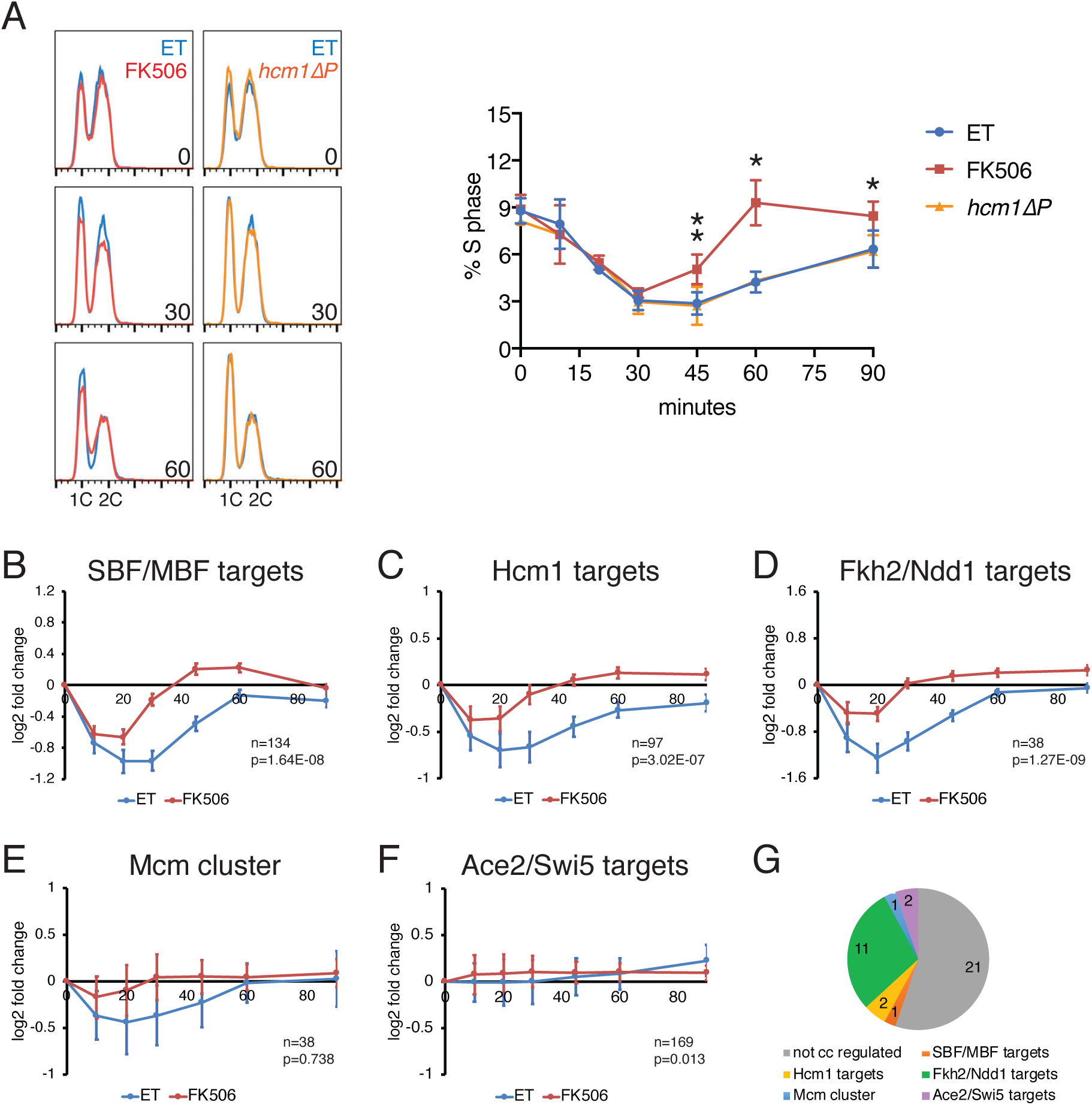
Regulation of the cell cycle by calcineurin. **(A)** *crz1Δor crz1Δhcm1ΔPcells* were treated with ET buffer (both gentoypes) or FK506 (*crz1Δ* only) for 15 minutes before the addition of 200 mM CaCl_2_. DNA content was measured by flow cytometry at the indicated time points. Overlays of FACS plots for selected time points are shown at the left. Quantitation of the percentage of cells in S phase in n=3 (ET, FK506) or n=2 *(hcm1ΔP)* experiments is shown on the right. Error bars represent standard deviations. Statistical significance between ET and FK506 samples was calculated for each time point using a paired t-test. Asterisks indicate *p<0.05, **p<0.01. Raw FACS plots for a representative experiment are shown in Figure S1A. **(B-F)** RNA-seq was performed on duplicate time course experiments with ET- and FK506-treated *crz1Δ* cells from (a). Shown are average expression of the indicated groups of cell cycle-regulated genes. Error bars indicate the 95% confidence interval. Number of genes (n) and the adjusted p-value indicating the significance of the difference between ET and FK506 curves are included. See Figure S2 for subsets of SBF/MBF and Ace2/Swi5 targets. **(G)** Genes whose expression is significantly different between ET and FK506-treated *crz1Δ* cells after 10 minutes of CaCl_2_ treatment. 17 of 38 genes (colored segments) are cell cycle regulated genes. See Table S1 for list of genes.

One mechanism by which CN could enforce a cell cycle arrest is by downregulating expression of genes that drive the cell cycle forward. The S-phase TF Hcm1 is an established target of CN (Arsenault et al., 2015); however, an Hcm1 mutant that cannot interact with CN had no effect on CN-mediated response to stress (*hcm1ΔP*, Figures 1A and S1A). To determine if CN impacts the activities of other cell cycle-regulatory TFs, RNA-seq was performed on samples from CaCl_2_ time course experiments and the average change in gene expression of targets of each TF was compared between control and FK506-treated cells. Interestingly, in addition to Hcm1 target genes (Figures 1C and S2A; Data S1), two additional clusters of genes were significantly downregulated by CN. These included targets of the SBF/MBF complexes (Ferrezuelo et al., 2010), which peak at the G1/S transition (Figures 1B and S2A-D, Data S1), as well as targets of the Fkh2/Ndd1 complex (also called the Clb2 cluster; Spellman et al., 1998), which peak at the G2/M transition (Figures 1D and S2A; Data S1). In contrast, although some genes in the Mcm cluster and Ace2/Swi5 clusters were also altered by FK506 treatment (Figures 1E, 1F, S2A, S2E and S2F; Data S1), neither cluster exhibited a coordinated response to CaCl_2_, suggesting that the activities of cell cycle TFs that control the expression of these genes may not be regulated by CN. Similar patterns of regulation were found when we re-analyzed a published dataset from CaCl_2_-treated wild-type cells (Yoshimoto et al., 2002) (Figures S1C-G), validating our findings. Importantly, CN-dependent downregulation of G1/S and G2/M genes was not an indirect consequence of a difference in cell cycle position, since the largest change in expression of these genes was at the 30-minute time point (Figures 1B and 1D), when ET-and FK506-treated cells had similar cell cycle distributions (Figure 1A). Thus, CN-dependent changes in cell cycle gene expression precede CN-dependent effects on cell cycle progression.

### CN activates the SAPK Hog1 to downregulate G1/S genes

We next sought to understand how CN downregulates the expression of G1/S genes. In budding yeast, the stress-activated MAPK Hog1 has two roles in controlling entry into S phase. Hog1 both inhibits the SBF and MBF TF complexes that drive expression of G1/S genes (Bellí et al., 2001; González-Novo et al., 2015), and increases levels of the Cdk1 inhibitors Sic1 and Cip1, which inhibit G1 cyclin/Cdk complexes and block entry into S phase (Chang et al., 2017; Escoté et al., 2004). Notably, many stresses that activate CN also increase osmolarity (Matsumoto et al., 2002). This raised the possibility that CaCl_2_ addition could promote both CN and Hog1 activation, and by extension lead to the downregulation of G1/S genes that we observed.

Consistent with our hypothesis, CaCl_2_ addition led to a rapid induction of Hog1 phosphorylation that persisted over the time course (Figures 2A and 2B). In addition, expression of genes that are known to be induced following Hog1 activation (Capaldi et al., 2008) mirrored the timing of Hog1 phosphorylation (Figure 2C). In particular, the well-characterized Hog1-responsive gene *STL1* was induced after 10 minutes of CaCl_2_ treatment in control cells and largely persisted throughout the time course (Figure 2D). Interestingly, this extended period of Hog1 activation in response to CaCl_2_ differed from the reported responses to other Hog1-activating stresses, which produce a transient peak of Hog1 activation that is rapidly reduced as homeostasis is restored (Brewster et al., 1993). Indeed, although Hog1 was phosphorylated to similar extents in response to CaCl_2_, NaCl, and sorbitol, Hog1 phosphorylation was quickly reversed when cells were continually exposed to NaCl or sorbitol, whereas it persisted in CaCl_2_-treated cells (Figure 2E). These results suggest that, in contrast to other osmostressors, CaCl_2_ generates an additional signal that prolongs Hog1 activation.

**Figure 2.**
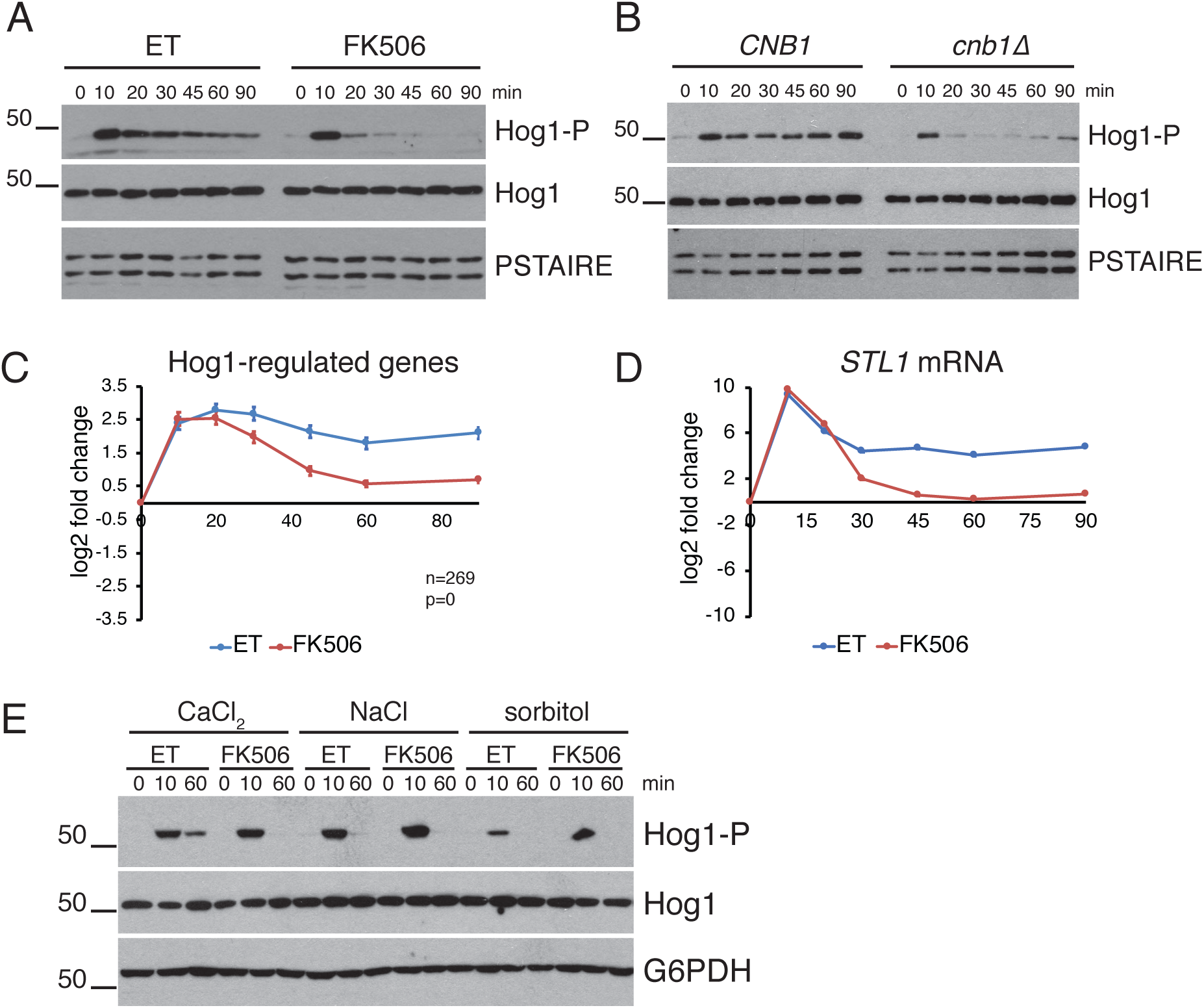
Calcineurin maintains Hog1 activation in CaCl_2_-treated cells. **(A)** Inhibition of CN with FK506 reduces Hog1 phosphorylation in response to CaCl_2_. *crz1Δ* cells were treated with ET buffer or FK506 for 15 minutes and then 200 mM CaCl_2_ was added. The activating phosphorylation on Hog1 (Hog1-P) was monitored by Western blot. Total Hog1 levels and PSTAIRE (loading control) are also shown. **(B)** Deletion of *CNB1* reduces Hog1 phosphorylation in response to CaCl_2_. *crz1Δ CNB1* and *crz1Δ cnb1Δ* cells were treated with 200mM CaCl_2_ and Hog1 activation was monitored by Western blot, as in (A). **(C)** Average expression of Hog1-regulated genes from RNA-seq experiments described in Fig 1. Number of genes (n) and the adjusted p-value indicating the significance of the difference between ET and FK506 curves are included. **(D)** RNA was collected from cells treated as in (A) and levels of *STΔ* mRNA were quantified by RT-qPCR. *STΔ* levels were normalized to *ACT1* and fold change calculated relative to the 0-minute time point. Log2 fold change from a representative experiment is shown. Error bars represent the standard deviation of technical replicates. Note that most error bars are too small to be visualized. **(E)** Comparison of Hog1-activating stressors. *crz1Δ* cells were pretreatedwith ET or FK506 for 15 minutes before the addition of 200 mM CaCl_2_, 400 mM NaCl or 1M sorbitol and samples were collected after the indicated number of minutes. Shown are Western blots for activated Hog1 (Hog1-P), total Hog1 and G6PDH (loading control).

Notably, Hog1 was phosphorylated in both FK506-treated cells and cells lacking the regulatory subunit of CN, Cnb1 (Figure 2A and 2B). However, initial CaCl_2_-induced Hog1 phosphorylation quickly decreased in the absence of CN activity, indicating that CN is required to maintain Hog1 phosphorylation in response to CaCl_2_ stress. Expression of Hog1-regulated genes mirrored the pattern of Hog1 phosphorylation, quickly returning to near starting levels when CN was inhibited, despite being induced to similar levels initially in control and FK506-treated cells (Figure 2C and 2D). A similar effect of CN on Hog1 activation was observed in a wild type (*CRZ1* proficient) strain background (Figures S3A and S3B), although cells adapted to the stress more quickly, as expected. Together, these data demonstrate that CN is required to maintain Hog1 activation in response to continuous CaCl_2_ exposure.

Since Hog1 is known to inhibit the G1/S transition, and it is induced equally by CaCl_2_ in control and FK506-treated cells, this suggested that Hog1 activation might promote the initial CN-independent cell cycle arrest that occurs in response to stress (Figure 1A). To test this possibility, time course experiments were performed in cells carrying a deletion of *HOG1*. Interestingly, while the fraction of Hog1-expressing cells in S phase decreased by 30 minutes after the addition of CaCl_2_ (Figure 1A), *hog1Δ* cells showed no significant change in cell cycle distribution after 30 minutes (Figure 3A and S4). However, by the 45-minute time point there was a CN-dependent decrease in the fraction of *hog1Δ* cells in S-phase (Figure 3A), similar to the decrease observed in cells with wild-type *HOG1* (Figure 1A). These data confirm that the initial CaCl_2_-induced arrest is the result of Hog1 activation, and suggest that CN maintains cell cycle arrest over time, through a mechanism that is at least partly independent of Hog1.

**Figure 3.**
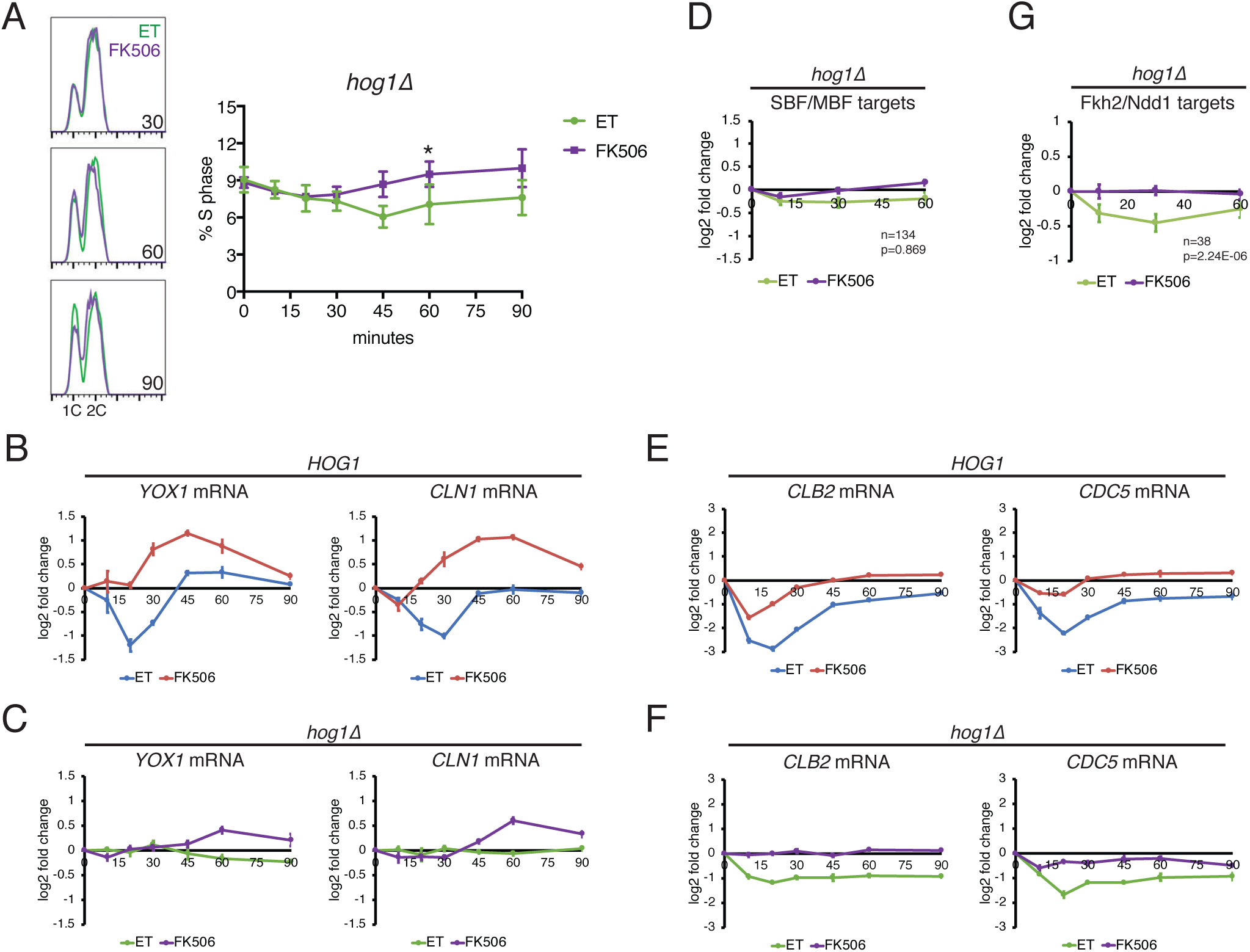
CN-dependent downregulation of G1/S genes requires Hog1. **(A)** *crz1Δ hog1Δ* cells were treated with ET buffer or FK506 for 15 minutes before the addition of 200 mM CaCl_2_. DNA content was measured by flow cytometry at the indicated time points. Overlays of FACS plots for selected time points are shown at the left. Quantitation of the percentage of cells in S phase in n=3 experiments is shown on the right. Error bars represent standard deviations. Statistical significance between ET and FK506 samples were calculated for each time point using a paired t-test. Asterisks indicate *p<0.05. Raw FACS plots for a representative experiment are included in Figure S4. **(B)** RT-qPCR of SBF/MBF target genes *YOX1* and *CLN1* following the addition of 200 mM CaCl_2_ to *crz1Δ* cells. Cells were pretreated with ET buffer or FK506, as indicated. Levels were normalized to *ACT1* and fold change calculated relative to the 0-minute time point. Log2 fold change from a representative experiment is shown. Error bars represent standard deviation of technical replicates. **(C)** RT-qPCRof *YOX1* and *CLN1*, as in (A), except experiment was performed in *crz1Δ hog1Δ* cells. **(D)** Average expression of all SBF/MBF target genes in *crz1Δ hog1Δ* cells at the indicated time points after CaCl_2_ addition, as measured by RNA-seq. Error bars indicate the 95% confidence interval. Number of genes (n) and the adjusted p-value indicating the significance of the difference between ET and FK506 curves are included. **(E)** RT-qPCR of Clb2 cluster genes *CLB2* and *CDC5* in *crz1Δ* cells, as in (A). **(F)** RT-qPCR of *CLB2* and *CDC5* in *crz1Δ hog1Δ* cells, as in (B). **(G)** Average expression of all Fkh2/Ndd1 target genes in *crz1Δ hog1Δ* cells, as measured by RNA-seq, as in (C).

We next tested whether Hog1 is required for CN-dependent downregulation of SBF/MBF target genes by measuring changes in cell cycle transcripts using RT-qPCR. First, we validated expression of two SBF/MBF target genes, *YOX1* and *CLN1*, which decreased in expression in RNA-seq experiments. Both genes were downregulated within 20 minutes after the addition of CaCl_2_, and downregulation was blocked in FK506-treated cells (Figure 3B). In addition, expression of both genes was upregulated above starting levels at later time points in FK506-treated cells, similar to the average expression of all SBF/MBF target genes (Figure 1B). In contrast, CaCl_2_ did not trigger these changes in *CLN1* and *YOX1* expression in *hog1Δ* cells (Figure 3C), indicating that CN-dependent changes in G1/S gene expression require Hog1. RNA-seq analysis revealed that this pattern was consistent among the entire cluster of SBF/MBF target genes: average expression of SBF/MBF target genes did not change in *hog1Δ* cells following CaCl_2_ addition, and expression was not significantly different between control and FK506-treated cells (Figure 3D). Therefore, we conclude that CN regulates SBF/MBF target genes indirectly, by promoting the activation of Hog1.

We also examined whether activation of Hog1 impacted expression of Fkh2/Ndd1 target genes that peak in G2/M phase, which were also downregulated upon CaCl_2_ stress (Figure 1D). In contrast to SBF/MBF target genes, the G2/M genes *CLB2* and *CDC5*, as well as all G2/M genes in aggregate (Figure 3E-G), exhibited CN-dependent downregulation in the absence of Hog1, although they were downregulated to a lesser extent than in Hog1-expressing cells. These results suggest that, unlike G1/S genes, Hog1-independent mechanisms contribute to downregulation of G2/M genes.

### CN regulates levels and phosphorylation of G2/M TFs

The cell cycle regulatory TF network operates as an oscillator, with activators in each cell cycle stage inducing expression of downstream TFs in the network (Horak et al., 2002; Orlando et al., 2008). The S-phase TF Hcm1 is part of this oscillatory network and it promotes expression of the downstream TFs Fkh2 and Ndd1 that activate genes at the G2/M transition (Pramila et al., 2006). Since we found that Fkh2/Ndd1 target genes were downregulated by CN, this raised the possibility that Hcm1 inactivation by CN leads to a failure to properly induce *FKH2* and *NDD1* expression, causing a subsequent reduction in expression of Fkh2/Ndd1 target genes. However, Fkh2/Ndd1 target genes were among the earliest genes downregulated in a CN-dependent manner (Figure 1G; Table S1). Following 10 minutes of CaCl_2_ treatment, 38 genes were significantly different between control and FK506-treated cells and 11 of these genes were Fkh2/Ndd1 targets, all of which were downregulated. This rapid decrease in expression suggests that downregulation of Fkh2/Ndd1 target genes does not depend upon decreased transcription of *FKH2* and *NDD1*, since the effects of reduced expression of the TFs on their target genes would likely require longer than 10 minutes. To test this possibility directly, levels of Fkh2/Ndd1 target genes were examined in cells expressing an Hcm1 mutant that lacks the CN-docking site and cannot be inactivated by CN (Arsenault et al., 2015). Notably, Fkh2/Ndd1 targets were downregulated nearly identically in *hcm1ΔP* cells and ET-treated control cells (Figure S2A). This result confirms that the activity of Fkh2/Ndd1 is not indirectly inactivated via CN targeting of Hcm1 and suggests that CN regulates these TFs through an independent mechanism.

To elucidate the mechanism of downregulation Fkh2/Ndd1 target genes, we examined the levels of all known G2/M regulatory TF proteins over time following CaCl_2_ addition to control and FK506-treated cells. G2/M genes could be downregulated if expression of an activating TF is decreased, or if expression of a repressive TF is increased. Among the TFs that activate G2/M gene transcription, Ndd1 was strongly downregulated in response to CaCl_2_, and this response was largely, but not completely blocked when CN was inhibited (Figure 4A). Ndd1 protein levels mirrored its mRNA levels (Figure 4B), and Ndd1 protein exhibited a short half-life that is unchanged in response to CaCl_2_ (Figure 4C), suggesting that downregulation of Ndd1 protein results from a CN-dependent decrease in its transcription. Fkh2 protein also decreased, although to a lesser extent, after 45 minutes of CaCl_2_ treatment. Although *FKH2* mRNA was strongly downregulated at the 10-minute time point (Figure 4B), the more modest change in Fkh2 protein levels is likely explained by the fact that the protein is more stable than Ndd1 (Figure 4C). A second potential mechanism for downregulation of G2/M genes could be increased expression of a transcriptional repressor. However, levels of the G2/M repressive TF Yox1 and its paralog Yhp1 decreased in response to CaCl_2_, consistent with the fact that transcription of these TFs decreases in response to CN activation (Figure S5). Together these data suggest that downregulation of G2/M genes results in part from decreased expression of the activating TFs Ndd1 and Fkh2.

**Figure 4.**
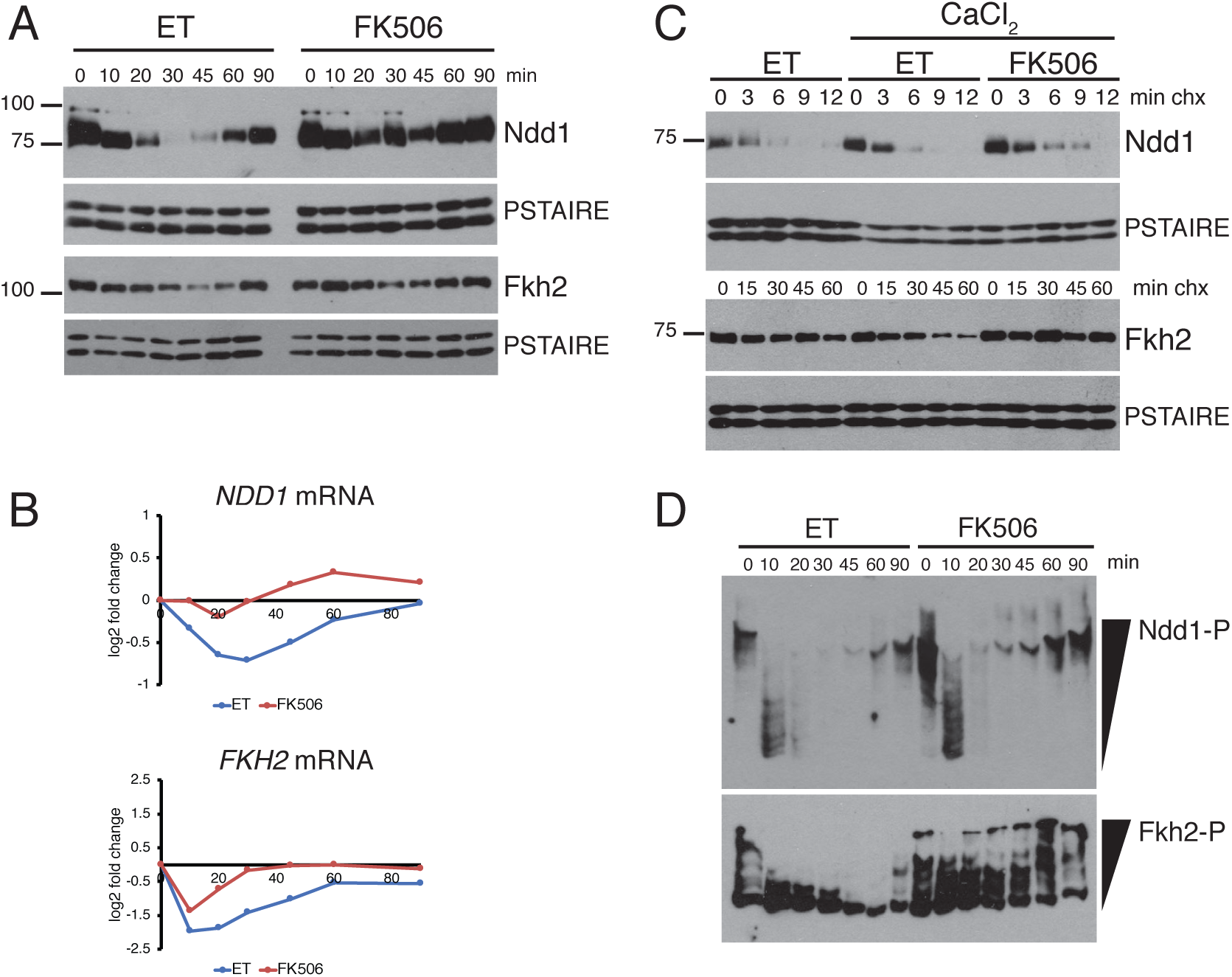
Regulation of G2/M TFs by CN. **(A)** Expression of TF proteins in response to CaCl_2_. Strains expressing the indicated tagged TFs were pretreated with ET buffer or FK506 for 15 minutes before the addition of 200mM CaCl_2_. Samples were collected for Western blotting at the indicated time points. Western blots were performed for a 3V5 tag on Ndd1 or a 3FLAG tag on Fkh2. PSTAIRE blots are shown as loading controls. **(B)** Expression of TF mRNAs in response to CaCl_2_. Shown are log2 fold change values, compared to the 0-minute time point, from RNA-seq experiments described in Figure 1. **(C)** Cycloheximide-chase assays of Ndd1 and Fkh2. Cells expressing 3V5-tagged Ndd1 and 3FLAG-tagged Fkh2 were pretreated with ET buffer or FK506 for 15 minutes, 200mM CaCl_2_ was added for an additional 2 minutes, then cycloheximide was added (0 minutes) and samples collected at the indicated time points for Western blot. **(D)** Phos-tag gel analysis of Fkh2 and Ndd1. ET and FK506 treated samples from Ndd1 and Fkh2 experiments in (A) were run on Phos-tag gels to visualize dephosphorylation in response to CaCl_2_ addition.

Our data indicated that Fkh2/Ndd1 target genes were downregulated as early as 10 minutes following the addition of CaCl_2_ (Figures 1D and 1G), although levels of Ndd1 and Fkh2 did not begin to decrease notably until later time points (Figure 4A). Therefore, other modes of regulation must also contribute to Fkh2/Ndd1 inactivation at the earliest time points after CaCl_2_ addition. Cdk1-mediated phosphorylation of both Fkh2 and Ndd1 is required for the recruitment of Ndd1 to promoters (Darieva et al., 2003; Pic-Taylor et al., 2004; Reynolds, 2003), so we examined whether either protein was dephosphorylated in response to CaCl_2_. Notably, both proteins were rapidly dephosphorylated following CaCl_2_ exposure (Figure 4D). Ndd1 was dephosphorylated in both control and FK506-treated cells, although the phosphorylated protein reappeared by the 20-minute time point when CN was inhibited. In contrast, Fkh2 dephosphorylation was blocked when CN was inhibited by FK506. Taken together, these data suggest that G2/M genes are initially downregulated as a result of dephosphorylation of Ndd1 and/or Fkh2, and this downregulation is enforced at later time points by a decrease in TF expression.

### CN regulates G2/M TFs through Hog1-dependent and -independent pathways

We next sought to disrupt the dephosphorylation and downregulation of Ndd1 and Fkh2 in response to CaCl_2_, to test whether regulation of these TFs is required for downregulation of G2/M genes. To accomplish this, we first investigated how phosphorylation of Fkh2 and Ndd1 is regulated in response to CaCl_2_. Two non-mutually exclusive possibilities were considered: CN could directly dephosphorylate Fkh2 or Ndd1, or alternatively, Cdk1 activity could be inhibited and this could lead to a decrease in phosphorylation of the TFs. Ndd1 is unlikely to be a direct CN substrate, since it was dephosphorylated similarly in control and FK506-treated cells (Figure 4D). In contrast, Fkh2 dephosphorylation was blocked by FK506. In addition, Fkh2 contains a PAISIS motif that matches the conserved CN docking site sequence and is in an accessible region of the protein (Goldman et al., 2014), making it a good candidate substrate of CN. However, we found that deletion of the predicted CN docking site in Fkh2 did not prevent its dephosphorylation *in vivo*, and we could not demonstrate dephosphorylation of Fkh2 upon treatment with CN *in vitro*, suggesting that Fkh2 regulation by CN is indirect (data not shown).

We considered whether CN might inhibit Cdk1 activity to promote dephosphorylation of G2/M TFs. In fact, CN was previously found to inhibit Clb2/Cdk1 activity indirectly, through its ability to dephosphorylate Hsl1 (Goldman et al., 2014; Mizunuma et al., 2001). Hsl1 is an inhibitor of the kinase Swe1 (Wee1 in other systems), which phosphorylates Cdk1 on Y19 to inhibit Cdk1 activity (Kellogg, 2003). Therefore, inactivation of Hsl1 by CN is thought to promote Swe1 activity, leading to Cdk1 inhibition in response to stress (Figure 5A). Consistent with this proposed role, we observed that levels of Y19-phosphorylated Cdk1 increased upon CaCl_2_ treatment, and this phosphorylation was completely eliminated upon deletion of *SWE1* (Figure 5B). We next tested whether the Hsl1-Swe1 pathway is required for dephosphorylation of Ndd1 and/or Fkh2 upon CN activation. Notably, dephosphorylation of both TFs was blocked in *swe1Δ* cells (Figure 5C), supporting the conclusion that it is inhibition of Cdk1 kinase activity that leads to their dephosphorylation. Importantly, deletion of *SWE1* also largely blocked the downregulation of Ndd1/Fkh2 target genes *CLB2, CDC5* and *CDC20* (Fig 5d), suggesting that dephosphorylation of Ndd1 and Fkh2 contributes to downregulation of G2/M genes.

**Figure 5.**
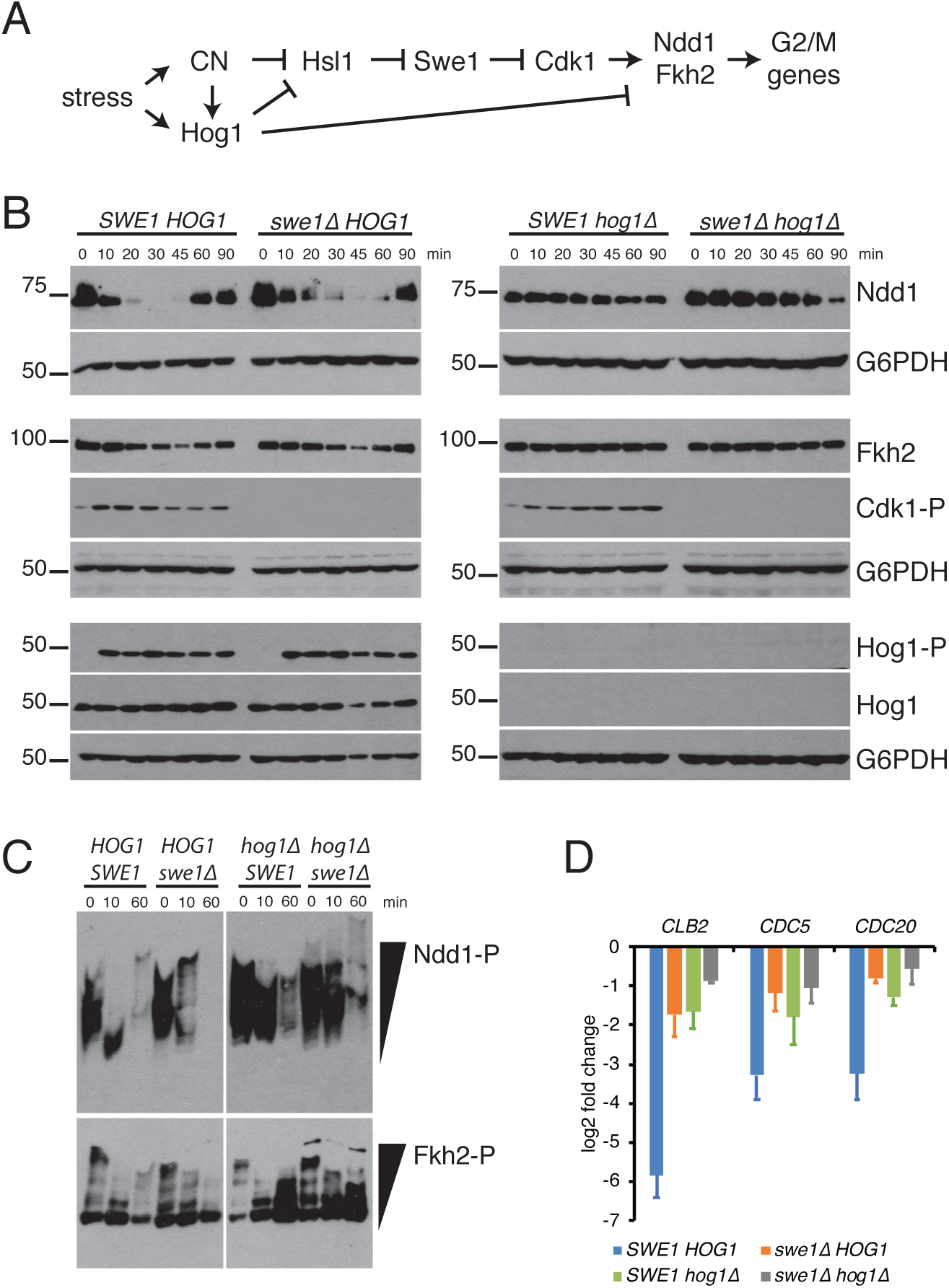
Swe1 and Hog1 regulate G2/M TF phosphorylation and activity. **(A)** Model of G2/M gene regulation in response to CaCl_2_ stress. **(B)** crz*1Δ*.dcells with the indicated mutations were treated with 200mM CaCl_2_ and cells were collected at the indicated time points for Western blot. Samples were assayed with antibodies against 3V5-tagged Ndd1, 3FLAG-tagged Fkh2, Cdk1 phosphorylated on Y19 (Cdk1-P), phosphorylated Hog1 (Hog1-P), total Hog1, and G6PDH (loading control). **(C)** Regulation of Ndd1 and Fkh2 phosphorylation. Phos-tag gels comparing phosphorylation of 3V5-tagged Ndd1 and 3FLAG-tagged Fkh2 in cells of the indicated genotypes after the addition of CaCl_2_ as in (A). Note that left and right halves of Ndd1 and Fkh2 panels come from the same gel, however different exposures are shown. **(D)** RT-qPCR of representative Fkh2/Ndd1 target genes 20 minutes after the addition of CaCl_2_ in cells with the indicated genotypes. Data is represented as log2 fold change compared to mRNA levels before the addition of CaCl_2_. Shown is an average of n=3 experiments, error bars represent standard deviations.

Interestingly, Hog1 has also been shown to inactivate Hsl1 to stimulate Swe1 activity (Clotet et al., 2006). In addition, we observed that downregulation of Fkh2/Ndd1 target genes was partially blocked in *hog1Δ* cells (Figures 3F and 3G). These observations suggested that Hog1 may also contribute to Cdk1 inhibition and the dephosphorylation of Ndd1 or Fkh2. In support of this possibility, deletion of *HOG1* reduced Cdk1 phosphorylation at early time points after the addition of stress (Figure 5B), and dephosphorylation of Ndd1 was largely blocked (Figure 5C). In contrast, Fkh2 dephosphorylation was unaffected in *hog1Δ* single mutant cells. However, dephosphorylation of both TFs was almost completely blocked in *hog1Δ swe1Δ* cells (Figure 5C). The fact that the double mutant exhibited a more complete block to dephosphorylation than either single mutant is consistent with the fact that that Hog1 and CN can independently activate Swe1. This result suggests that CN promotes Ndd1/Fkh2 dephosphorylation in part through activation of Hog1 and in part through its ability to directly dephosphorylate and inactivate Hsl1 (Figure 5A).

Although our results indicate that Hog1 contributes to Cdk1 inhibition and the rapid dephosphorylation of Ndd1 in response to stress (Figure 5C), we also observed that Hog1 was required for the downregulation of Ndd1 and Fkh2 proteins that occurred at later time points (Figure 5A). In contrast to Hog1-expressing cells, neither protein decreased in levels when *hog1Δ* cells were exposed to stress (Figure 5B). Importantly, deletion of *SWE1* and *HOG1* together almost completely blocked both dephosphorylation and reduction of Ndd1 and Fkh2 levels. Together, these data demonstrate that Hog1 and Swe1 both play key roles downstream of CN to dephosphorylate and downregulate expression of G2/M TFs (Figure 5A).

Having established that CN-induced changes in Ndd1 and Fkh2 expression did not occur in cells lacking both Hog1 and Swe1, we next tested whether G2/M genes were downregulated in response to stress when these pathways were disrupted. Notably, deletion of either *SWE1* or *HOG1* largely prevented downregulation of the G2/M genes *CLB2, CDC5* and *CDC20* (Figure 5D). Furthermore, *swe1Δ hog1Δ* double delete cells showed a slightly stronger effect on the same targets. The stronger effect in *swe1Δ hog1Δ* cells relative to either single mutant to (i) block dephosphorylation of G2/M TFs and (ii) block downregulation of G2/M gene expression supports the conclusion that CN promotes G2/M TF inactivation through both Hog1-dependent and - independent mechanisms.

### CN arrests the cell cycle at multiple stages

Our data demonstrate that CN activation leads to widespread downregulation of cell cycle-regulated genes. We also found that CN contributes to cell cycle arrest in response to stress (Figure 1A). However, it is difficult to discern whether one or multiple phases of the cell cycle are blocked in response to CN activation when experiments are performed in asynchronous cultures. To determine whether progression through one or more phases is inhibited, we assayed the consequences of CN activation on cells synchronized in specific phases (Figure 6A). First, we examined how CaCl_2_ affects cells when they are released from a G1 arrest. In the absence of CaCl_2_, cells with and without the CN subunit Cnb1 released from a G1 arrest with similar kinetics (Figures 6B, green and purple, and S6A). However, when CN-expressing cells were released into the media containing CaCl_2_ there was a significant delay in progression through S phase, and this delay was largely reversed in *cnb1Δ* cells (Figures 6B, compare blue and red, and S6A). Similar results were obtained when CN was inhibited by FK506 (Figures 6C and S6B). This CN-dependent G1/S delay could be the result of cells arresting at the G1/S transition, or it could occur if cells proceed through the G1/S transition but CN inhibits DNA replication. To distinguish between these possibilities, cells were released from an early S-phase block (after the G1/S transition but prior to DNA replication; Figure 6A). Interestingly, although the addition of CaCl_2_ caused a modest delay in progression through S phase, there was no significant difference between control and FK506-treated cells (Figures 6D and S6C). Thus, CN activation acts on the G1/S transition to delay entry into S phase, consistent with its ability to inhibit expression of SBF/MBF target genes.

**Figure 6.**
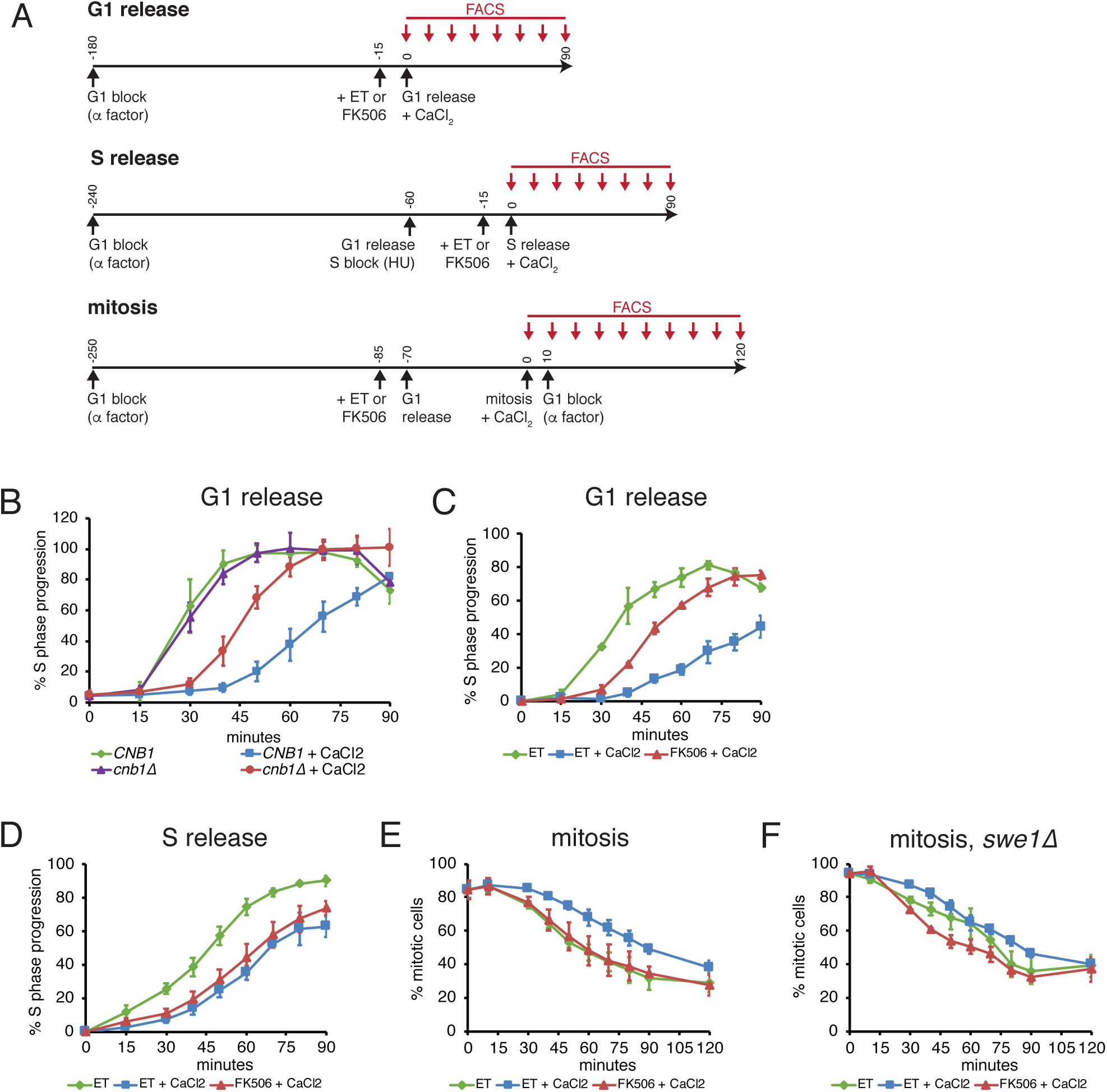
CN activation delays the cell cycle at multiple stages. **(A)** Schematic depicting different synchronization protocols used in subsequent panels. **(B-G)** Cells were synchronized before CaCl_2_ addition as indicated in individual panels and described in part (A). S phase progression and % mitotic cells were calculated as described in the Materials and Methods. In each panel, an average of n=3 (parts B, C, D and F) or n=4 (part E) is shown and error bars represent standard deviations. (b) *crz1Δ* cells with or without *CNB1* were released from a G1 arrest. (C) *crz1Δ* cells treated with ET buffer or FK506 were released from a G1 arrest. (D) *crz1Δ* cells were released from an S phase arrest. (E) *crz1Δ* cells were synchronized in mitosis. (F) *crz1Δ swe1Δ* cells were synchronized in mitosis.

CN also downregulates expression of G2/M genes, so we next tested whether CN delays progression through mitosis. Previous studies have implicated CN in arresting cells at G2/M phase through its ability to activate Swe1 (Mizunuma et al., 2001; 1998; Yokoyama et al., 2006). However, these experiments were performed in cells lacking a subunit of the PP2A phosphatase (Zds1), which results in elevated levels of Swe1 and a G2/M delay even in the absence of CN activation. Notably, we observed a CN-dependent mitotic delay when CaCl_2_ was added to Zds1-expressing cells that were synchronized in mitosis (Figures 6E and S6D). Moreover, this delay was almost entirely eliminated in *swe1Δ* cells (Figures 6F and S6E), consistent with the fact that Fkh2/Ndd1 target genes are not downregulated in this mutant (Figure 5C). Thus, Swe1 (and by extension its ability to inhibit Cdk1 activity) is required for CN-dependent downregulation of G2/M genes and the mitotic delay observed when cells receive CaCl_2_ stress. Together, these analyses support the conclusion that CN signals to the cell cycle regulatory network in multiple stages in order to delay cell cycle progression while cells adapt to environmental stress.

## Discussion

Many stress response pathways target cell cycle regulatory TFs that control key transitions. Since cell cycle-regulatory TFs are part of an oscillatory and interconnected network (Horak et al., 2002; Orlando et al., 2008), we set out to examine the effect of the stress-activated phosphatase CN on the entire network, and to follow changes over time as cells respond and adapt to CaCl_2_ stress. This time-resolved analysis revealed that CN has a broad role in rewiring cell cycle-regulated transcription and arresting the cell cycle. Although previous studies have identified a few direct CN substrates that impact the cell cycle (Arsenault et al., 2015; Goldman et al., 2014; Mizunuma et al., 2001), our findings suggest that many of the transcriptional and cell cycle changes that occur downstream of CN result from crosstalk to the stress-activated MAPK Hog1.

Our results present a dynamic picture of how CN and Hog1 collaborate to influence the cell cycle when cells are exposed to CaCl_2_. Initially, CaCl_2_ causes a change in osmolarity that activates Hog1, and the influx of Ca^2+^ ions into the cell activates CN. Changes in osmolarity are known to activate Hog1 within minutes, however in most cases this activation is rapidly shut off as glycerol is synthesized and homeostasis is restored (Muzzey et al., 2009). We show here that CaCl_2_ affects Hog1 differently than other osmostressors, because CN provides a signal that maintains levels of active Hog1 over time.

Together CN and Hog1 promote widespread changes in the expression of cell cycle-regulated genes (Figure 7). The concurrent activation of these pathways triggers an immediate downregulation of several clusters of cell cycle genes within 10 minutes of CaCl_2_ treatment: Hog1 mediates inactivation of SBF and MBF to downregulate G1/S genes (Bellí et al., 2001; González-Novo et al., 2015), CN dephosphorylates and inactivates the S-phase TF Hcm1 (Arsenault et al., 2015), and CN and Hog1 both contribute to Swe1 activation, which results in dephosphorylation of G2/M TFs Ndd1 and Fkh2. After approximately 20 minutes, inactivation of these TFs is reinforced as CN functions to maintain Hog1 activation, and levels of Fkh2 and Ndd1 proteins decrease through a Hog1-dependent pathway. Finally, after approximately 90 minutes of CaCl_2_ exposure, cells begin to adapt to the stress, levels and phosphorylation of TFs are returned to near starting levels, expression of cell cycle genes is restored, and cells resume cycling.

**Figure 7.**
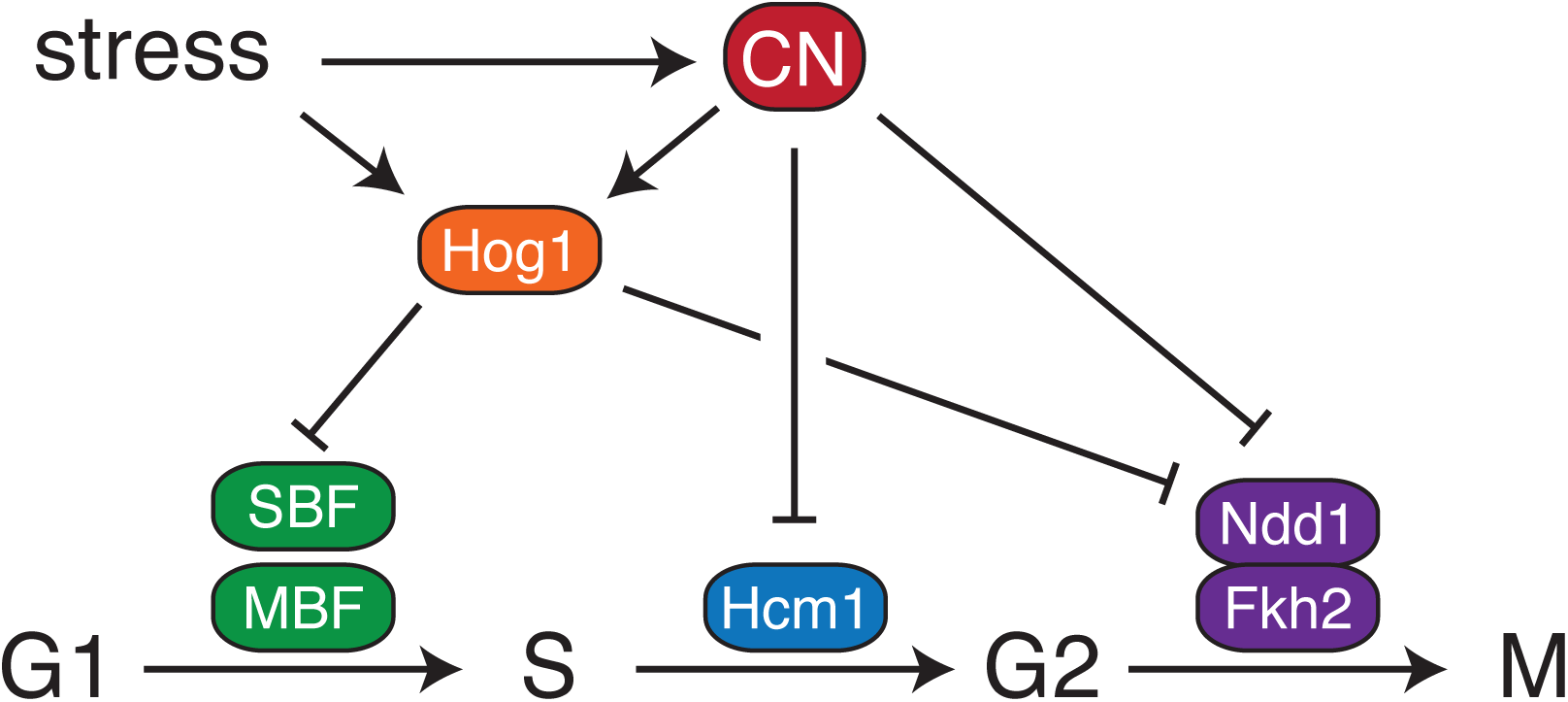
Calcineurin rewires cell cycle-regulated gene expression through multiple mechanisms. In response to stress, CN regulates different clusters of cell cycle genes through distinct mechanisms. G1/S genes are downregulated indirectly through the ability of CN to stimulate Hog1 activity, which in turn inactivates G1/S TFs. The S-phase TF Hem1 is directly inhibited by CN-mediated dephosphorylation (Arsenault et al., 2015). Finally, CN inactivates G2/M TFs through both Hog1-dependent and -independent mechanisms.

Since G2/M TFs Ndd1 and Fkh2 are Hcm1 target genes (Pramila et al., 2006), this suggested that inactivation of Hcm1 by CN (Arsenault et al., 2015) might promote downregulation of G2/M genes. However, we saw no significant change in downregulation of Fkh2/Ndd1 target genes in cells expressing a CN-resistant Hcm1 protein (Figure S2A). This result is consistent with the fact that inactivating a pathway by blocking transcription is a slow response, as it depends upon shutting off expression of the TFs and waiting for the TF proteins to be degraded before having an impact on their target genes. Rapid responses are more likely to result from modulating the phosphorylation landscape, which supports our finding that the timing of Fkh2 and Ndd1 dephosphorylation correlates well with the downregulation of their target genes (Figure 5). Although downregulation of Ndd1 and Fkh2 proteins likely contributes to a decrease in target gene expression after prolonged exposure to stress.

Surprisingly, neither Ndd1 nor Fkh2 appear to be direct substrates of the phosphatase CN, despite the fact that Fkh2 has features that suggest it might be a CN target (Goldman et al., 2014). Although our attempts to demonstrate dephosphorylation of Fkh2 by CN in vitro have been unsuccessful, it remains possible that CN could target a phosphosite that we cannot detect by a gel shift in our experiments. Despite this possibility we see that bulk Fkh2 dephosphorylation is significantly impaired in *hog1Δ swe1Δ* cells that are treated with CaCl_2_, supporting the model that dephosphorylation is indirectly regulated by Cdk1 inhibition.

CN not only inhibits expression of multiple clusters of cell cycle genes, but it also delays the cell cycle at multiple stages. Our findings are consistent with previous evidence suggesting that CN blocks G2/M progression through activation of Swe1. However, previous studies were carried out in a sensitized genetic background lacking the PP2A subunit Zds1 (Mizunuma et al., 1998), and we show that this arrest also happens in Zds1-expressing cells. We also show for the first time that CN mediates a transient G1/S arrest. Our data suggests that this G1/S arrest occurs through crosstalk to the Hog1 pathway, since downregulation of SBF/MBF target genes and initial cell cycle arrest do not occur in *hog1Δ* cells. This Hog1-dependent arrest likely occurs through a combination of its effect on transcription and its ability to upregulate Cdk1 inhibitors, as previously described (Bellí et al., 2001; Chang et al., 2017; Escoté et al., 2004; González-Novo et al., 2015).

Most environmental stresses trigger a common transcriptional response, termed the environmental stress response, which includes approximately 300 induced and 600 repressed genes (Gasch et al., 2000). However, each individual stress also activates a unique set of stress-specific changes in gene expression, which enable cells to adapt to the unique stressor. Interestingly, many environmental stressors simultaneously activate multiple signaling pathways, which may help cells tailor the specific transcriptional response to a particular stress. Hog1 and its mammalian homolog p38 in particular are activated by a wide array of stressors that also signal to other pathways including heat shock, oxidative stress, glucose starvation and arsenite (de Nadal et al., 2002; Ono and Han, 2000; Piao et al., 2012; Sotelo and Rodríguez-Gabriel, 2006; Thorsen et al., 2006; Winkler et al., 2002). CN is similarly activated by a number of stressors that stimulate multiple pathways. Cell wall stress activates CN in addition to the MAPK Mpk1/Slt2 (Levin, 2011). Additionally, many of the canonical CN-activating stresses are ions that cause hyperosmotic stress and also activate Hog1 (Farcasanu et al., 1995; Matsumoto et al., 2002). This overlap in the activation of stress response pathways makes it likely that the crosstalk between CN and Hog1 occurs in response to other stimuli in addition to CaCl_2_ stress.

Alternate stress response pathways have also been found to crosstalk to the Hog1 pathway. For example, the MAPK Mpk1 stimulates Hog1 activation in response to cell wall stress or heat shock by promoting loss of turgor pressure that activates the osmosensor Sln1 (Baltanas et al., 2013; Dunayevich et al., 2018). To our knowledge our findings are the first report of a stress-activated phosphatase that stimulates Hog1 activation. However, the cell cycle-regulatory phosphatase Cdc14 has been shown to contribute to Hog1-dependent downregulation of cell cycle genes in response to NaCl (Chasman et al., 2014). In contrast to what we find with CN, Cdc14 activity is required for Hog1 to be fully activated immediately after osmostress is applied, whereas CN is required to maintain Hog1 activation over time. Therefore, CN is likely to stimulate Hog1 activity through a distinct mechanism.

Although the best understood functions of CN in mammals are in regulation of the immune system (Hogan et al., 2003; Vaeth and Feske, 2018), recent evidence suggests it also plays a part in the stress response (Medina et al., 2015; Zhang et al., 2016). Oxidative stress and nutrient limitation both lead to the release of intracellular Ca^2+^ stores that activate CN. Upon activation, CN targets the transcription factor TFEB to induce expression of lysosomal biogenesis and autophagy genes. Notably, p38 is also activated by stress signals and similarly downregulates cell cycle-regulated gene expression and arrests the cell cycle in mammalian cells (Gubern et al., 2016). This suggests that the crosstalk between CN and Hog1 pathways may be conserved among eukaryotes to help cells tune their response to specific types of environmental stress.

## Methods

### Yeast strains and growth conditions

A complete list of strains is included in Table S2. Strains carrying gene deletions and epitope tags were constructed using standard methods (Longtine et al., 1998; Rothstein, 1991). All strains were grown in rich medium (YM-1) or synthetic complete medium (C) with 2% dextrose at 30°C (Benanti et al., 2007). In all experiments that include CaCl_2_ treatment, strains were grown in C medium with 1% ammonium chloride as the nitrogen source. Where indicated, cells were pretreated with ET buffer (90% ethanol, 10% Tween-20) or 1 μg/ml FK506 (LC Laboratories) in ET buffer 15 minutes before the addition of 200mM CaCl_2_. For experiments with NaCl or sorbitol treatment, cells were also grown in synthetic complete medium with ammonium chloride and pretreated with ET buffer or FK506 for 15 minutes before the addition of 400 mM NaCl or 1 M sorbitol.

### Flow cytometry

Cells were fixed in 70% ethanol and stained with Sytox Green (Invitrogen) as previously described (Landry et al., 2012). DNA content was measured on a FACScan (Becton Dickinson) or a Guava easyCyte HT (Millipore) flow cytometer. Data was analyzed using FlowJo (FlowJo, LLC) software. To quantify percent of cells in S-phase (Figures 1A and 3A), percentage of cells with a DNA content between 1C and 2C was calculated.

### RNA-seq and data analysis

Total RNA was isolated from 5 OD of cells and purified as previously described (Schmitt et al., 1990). Library preparation and sequencing was performed by BGI Genomic Services. Briefly, ribosomal RNAs were depleted and strand-specific libraries constructed prior to 50 base single-end sequencing on an Illumina HiSeq4000 platform. Two biological replicates of each time course were performed.

For all analyses, default parameters were used unless otherwise specified. Raw reads from RNA-seq experiments were assessed for their quality using fastqc (http://www.bioinformatics.bbsrc.ac.uk/projects/fastqc), followed by alignment to the *Saccharomyces cerevisiae* reference genome R64-1-1 from Ensembl using Tophat2 (version 2.1.1) (Kim et al., 2013) with the maximal intron length set as 100 kb. HTseq (Anders et al., 2015) was used to generate a gene-level count table, which was subsequently used for differential gene expression analysis using the voom method implemented in the limma package (version 3.32.10) (Law et al., 2014). Correlation within the same condition and genotype was estimated using the *duplicateCorrelation* function from the limma package.

A linear model was fit to include two independent variables, i.e., experimental condition (a combination of genotype, treatment and time post treatment) and batch, and with the correlation estimated above as input to the lmFit function. Tests of the following predefined contrasts were performed within the linear model framework. Gene expression at each time point post treatment was compared to its corresponding 0-minute baseline within the same treatment and genotype. In addition, changes of gene expression from baseline to each time point were compared between ET and FK506 treatments. Genes with a BH-adjusted p-value (Benjamini and Hochberg, 1995) less than 0.05 were considered as significantly differentially expressed for each comparison. Distance-based differential analyses of gene curves were performed as described (Minas et al., 2011) to compare CaCl_2_ time course data between ET and FK506-treated cells.

### Western blots

To prepare samples for Western blotting, equivalent optical densities of cells were collected and lysed in cold TCA buffer (10 mM Tris pH 8.0, 10% trichloroacetic acid, 25 mM ammonium acetate, 1 mM EDTA). After incubation on ice, lysates were centrifuged and pellets resuspended in resuspension solution (100 mM Tris pH 11.0, 3% SDS). Samples were heated to 95°C for 5 minutes and clarified by centrifugation. 4X SDS-PAGE sample buffer (250 mM Tris pH 6.8, 8% SDS, 40% glycerol, 20% β-mercaptoethanol) was added to clarified lysates before heating to 95°C for an additional 5 minutes. Western blotting was performed with antibodies against Hog1 (sc-6815 or sc-165978, Santa Cruz Biotechnology), phosphorylated Hog1 (anti-Phospho-p38 MAPK, 9211L, Cell Signaling Technology), PSTAIRE (sc-53, Santa Cruz Biotechnology or P7962, Sigma), G6PDH (A9521, Sigma), V5 (R96025, Invitrogen), FLAG (clone M2, F1804, Sigma), MYC (clone 9E10, M5546, Sigma), Clb2 (sc-9071, Santa Cruz Biotechnology), or Y19-phosphorylated Cdk1 (anti-Phospho-cdc2, 9111S, Cell Signaling Technology).

For Phos-tag gels, resolving gels contained 6% acrylamide/bis solution 29:1, 386 mM Tris-Cl, pH 8.8, 0.1% SDS, 0.2% ammonium persulfate (APS), 25 μM Phos-tag acrylamide (Wako), 50 μM manganese chloride, and 0.17% tetramethylethylenediamine (TEMED) and stacking gels contained 5% acrylamide/bis solution 37.5:1, 126 mM Tris-Cl, pH 6.8, 0.1% SDS, 0.1% APS, and 0.04% TEMED. Gels were washed twice in 100 ml of transfer buffer containing 10 mM EDTA for 15 minutes and once in 50 ml of transfer buffer for 10 minutes before transferring to nitrocellulose.

### RT-qPCR

Reverse transcription was carried out with 2µg of total RNA and random primers (Promega), followed by treatment with RNase H (New England Biolabs). Quantitative PCR was preformed using 2X SYBR Fast Master Mix Universal (Kapa Biosystems) and primers for the indicated genes (see Table S3 for primer sequences) using a Mastercycler EP Realplex thermocycler (Eppendorf). mRNA levels were normalized to *ACT1* and fold change values were calculated by comparing the normalized expression at the indicated time to the expression of the target before CaCl_2_ addition.

### Cycloheximide-chase assays

Cells were treated with ET buffer or FK506 for 15 minutes before the addition of 200 mM CaCl_2_. Two minutes after CaCl_2_ addition, 50 µg/ml cycloheximide (Biomatik) was added to block protein synthesis. Samples were collected after the indicated number of minutes for Western blot analysis.

### Cell cycle synchronization

For G1 arrest-release experiments, cells were treated with 10 µg/ml alpha factor for three hours (with an additional equivalent amount re-added after 2 hours) and then released into media without alpha factor and with or without 200 mM CaCl_2_. 15 minutes prior to release, cells were treated with ET buffer or FK506 as indicated. To synchronize cells in S phase, cells were release from a three-hour alpha factor arrest into medium containing 200 mM hydroxyurea (HU) for an additional hour before releasing into medium without HU and with or without 200 mM CaCl_2_. To synchronize cells in mitosis, cells were arrested in G1 with alpha factor for 3 hours, with ET or FK506 added after 2 hours and 45 minutes. Cells were then released into media without alpha factor (containing ET/FK506) and CaCl_2_ was added 70 minutes after release from the arrest, when greater than 80% of cells had a 2C DNA content. Alpha-factor was added back 10 minutes after the addition of CaCl_2_, to block cells in the subsequent G1 phase. S-phase progression and percentage of mitotic cells were calculated from the mean of the DNA content in each population, as previously described (Willis and Rhind, 2009). Data are presented as average values from a minimum of three experiments with error bars representing standard deviations.

## Acknowledgements

The authors thank Tom Fazzio, Martha Cyert, Michelle Conti, Michael Lee and Marian Walhout for valuable discussions and critical reading of the manuscript. This work was supported by grant R01GM117152 from the National Institutes of Health to J.A.B.

## Author Contributions

C.M.L, M.J.F and J.A.B. conceived and designed the experiments; C.M.L., M.J.F. and H.E.A. performed the experiments; C.M.L., M.J.F., H.E.A., J.O., H.L., L.J.Z and J.A.B. analyzed the data; J.A.B. wrote the paper with input from all authors.

## Supplemental Information

**Data S1.** Changes in cell cycle-regulated gene expression in response to CaCl_2_ stress.

**Data S2.** Changes in cell cycle-regulated gene expression in response to CaCl_2_ stress in *hog1Δ* cells.

**Table S1.** CN-dependent changes in gene expression after 10 minutes of CaCl_2_ stress.

**Table S2.** Strain table.

**Table S3.** Primer table.

## References

Anders, S., Pyl, P.T., Huber, W., 2015. HTSeq--a Python framework to work with high-throughput sequencing data. Bioinformatics 31, 166–169. doi:10.1093/bioinformatics/btu638

Arsenault, H.E., Roy, J., Mapa, C.E., Cyert, M.S., Benanti, J.A., 2015. Hcm1 integrates signals from Cdk1 and Calcineurin to control cell proliferation. Mol Biol Cell 26, 3570–3577. doi:10.1091/mbc.E15-07-0469

Baltanas, R., Bush, A., Couto, A., Durrieu, L., Hohmann, S., Colman-Lerner, A., 2013. Pheromone-induced morphogenesis improves osmoadaptation capacity by activating the HOG MAPK pathway. Sci Signal 6, ra26–ra26. doi:10.1126/scisignal.2003312

Bellí, G., Gari, E., Aldea, M., Herrero, E., 2001. Osmotic stress causes a G1 cell cycle delay and downregulation of Cln3/Cdc28 activity in Saccharomyces cerevisiae. Mol Microbiol 39, 1022–1035.

Benanti, J.A., Cheung, S.K., Brady, M.C., Toczyski, D.P., 2007. A proteomic screen reveals SCFGrr1 targets that regulate the glycolytic-gluconeogenic switch. Nat Cell Biol 9, 1184–1191. doi:10.1038/ncb1639

Benjamini, Y., Hochberg, Y., 1995. Controlling the False Discovery Rate: A Practical and Powerful Approach to Multiple Testing. Journal of the Royal Statistical Society: Series B (Methodological) 57, 289–300. doi:10.1111/j.2517-6161.1995.tb02031.x

Bertoli, C., Skotheim, J.M., de Bruin, R.A.M., 2013. Control of cell cycle transcription during G1 and S phases. Nat Rev Mol Cell Biol 14, 518–528. doi:10.1038/nrm3629

Brewster, J.L., de Valoir, T., Dwyer, N.D., Winter, E., Gustin, M.C., 1993. An osmosensing signal transduction pathway in yeast. Science 259, 1760–1763.

Capaldi, A.P., Kaplan, T., Liu, Y., Habib, N., Regev, A., Friedman, N., O’Shea, E.K., 2008. Structure and function of a transcriptional network activated by the MAPK Hog1. Nat Genet 40, 1300–1306. doi:10.1038/ng.235

Chang, Y.-L., Tseng, S.-F., Huang, Y.-C., Shen, Z.-J., Hsu, P.-H., Hsieh, M.-H., Yang, C.-W., Tognetti, S., Canal, B., Subirana, L., Wang, C.-W., Chen, H.-T., Lin, C.-Y., Posas, F., Teng, S.-C., 2017. Yeast Cip1 is activated by environmental stress to inhibit Cdk1–G1 cyclins via Mcm1 and Msn2/4. Nature Communications 8, 1–13. doi:10.1038/s41467-017-00080-y

Chasman, D., Ho, Y.-H., Berry, D.B., Nemec, C.M., MacGilvray, M.E., Hose, J., Merrill, A.E., Lee, M.V., Will, J.L., Coon, J.J., Ansari, A.Z., Craven, M., Gasch, A.P., 2014. Pathway connectivity and signaling coordination in the yeast stress-activated signaling network. Mol. Syst. Biol. 10, 759–759. doi:10.15252/msb.20145120

Clotet, J., Escoté, X., Adrover, M.A., Yaakov, G., Garí, E., Aldea, M., de Nadal, E., Posas, F., 2006. Phosphorylation of Hsl1 by Hog1 leads to a G2 arrest essential for cell survival at high osmolarity. EMBO J 25, 2338–2346. doi:10.1038/sj.emboj.7601095

Cyert, M.S., Philpott, C.C., 2013. Regulation of cation balance in Saccharomyces cerevisiae. Genetics 193, 677–713. doi:10.1534/genetics.112.147207

Darieva, Z., Pic-Taylor, A., Boros, J., Spanos, A., Geymonat, M., Reece, R.J., Sedgwick, S.G., Sharrocks, A.D., Morgan, B.A., 2003. Cell cycle-regulated transcription through the FHA domain of Fkh2p and the coactivator Ndd1p. Current Biology 13, 1740–1745.

de Nadal, E., Alepuz, P.M., Posas, F., 2002. Dealing with osmostress through MAP kinase activation. EMBO Rep. 3, 735–740. doi:10.1093/embo-reports/kvf158

Dunayevich, P., Baltanas, R., Clemente, J.A., Couto, A., Sapochnik, D., Vasen, G., Colman-Lerner, A., 2018. Heat-stress triggers MAPK crosstalk to turn on the hyperosmotic response pathway. Sci Rep 8, 15168. doi:10.1038/s41598-018-33203-6

Escoté, X., Zapater, M., Clotet, J., Posas, F., 2004. Hog1 mediates cell-cycle arrest in G1 phase by the dual targeting of Sic1. Nat Cell Biol 6, 997–1002. doi:10.1038/ncb1174

Farcasanu, I.C., Hirata, D., Tsuchiya, E., Nishiyama, F., Miyakawa, T., 1995. Protein phosphatase 2B of Saccharomyces cerevisiae is required for tolerance to manganese, in blocking the entry of ions into the cells. Eur. J. Biochem. 232, 712–717.

Ferrezuelo, F., Colomina, N., Futcher, B., Aldea, M., 2010. The transcriptional network activated by Cln3 cyclin at the G1-to-S transition of the yeast cell cycle. Genome Biol 11, R67. doi:10.1186/gb-2010-11-6-r67

Gasch, A.P., Spellman, P.T., Kao, C.M., Carmel-Harel, O., Eisen, M.B., Storz, G., Botstein, D., Brown, P.O., 2000. Genomic expression programs in the response of yeast cells to environmental changes. Mol Biol Cell 11, 4241–4257.

Goldman, A., Roy, J., Bodenmiller, B., Wanka, S., Landry, C.R., Aebersold, R., Cyert, M.S., 2014. The Calcineurin Signaling Network Evolves via Conserved Kinase-Phosphatase Modules that Transcend Substrate Identity. Mol Cell. doi:10.1016/j.molcel.2014.05.012

González-Novo, A., Jiménez, J., Clotet, J., Nadal-Ribelles, M., Cavero, S., de Nadal, E., Posas, F., 2015. Hog1 targets Whi5 and Msa1 transcription factors to downregulate cyclin expression upon stress. Mol Cell Biol 35, 1606–1618. doi:10.1128/MCB.01279-14

Gubern, A., Joaquin, M., Marquès, M., Maseres, P., Garcia-Garcia, J., Amat, R., González-Nuñez, D., Oliva, B., Real, F.X., de Nadal, E., Posas, F., 2016. The N-Terminal Phosphorylation of RB by p38 Bypasses Its Inactivation by CDKs and Prevents Proliferation in Cancer Cells. Mol Cell 64, 25–36. doi:10.1016/j.molcel.2016.08.015

Haase, S.B., Wittenberg, C., 2014. Topology and Control of the Cell-Cycle-Regulated Transcriptional Circuitry. Genetics 196, 65–90. doi:10.1534/genetics.113.152595

Ho, Y.-H., Gasch, A.P., 2015. Exploiting the yeast stress-activated signaling network to inform on stress biology and disease signaling. Curr Genet 61, 503–511. doi:10.1007/s00294-015-0491-0

Hogan, P.G., Chen, L., Nardone, J., Rao, A., 2003. Transcriptional regulation by calcium, calcineurin, and NFAT. Genes & Development 17, 2205–2232. doi:10.1101/gad.1102703

Horak, C.E., Luscombe, N.M., Qian, J., Bertone, P., Piccirrillo, S., Gerstein, M., Snyder, M., 2002. Complex transcriptional circuitry at the G1/S transition in Saccharomyces cerevisiae. Genes & Development 16, 3017–3033. doi:10.1101/gad.1039602

Kellogg, D.R., 2003. Wee1-dependent mechanisms required for coordination of cell growth and cell division. J Cell Sci 116, 4883–4890. doi:10.1242/jcs.00908

Kim, D., Pertea, G., Trapnell, C., Pimentel, H., Kelley, R., Salzberg, S.L., 2013. TopHat2: accurate alignment of transcriptomes in the presence of insertions, deletions and gene fusions. Genome Biol 14, R36. doi:10.1186/gb-2013-14-4-r36

Landry, B.D., Doyle, J.P., Toczyski, D.P., Benanti, J.A., 2012. F-Box Protein Specificity for G1 Cyclins Is Dictated by Subcellular Localization. PLoS Genet 8, e1002851. doi:10.1371/journal.pgen.1002851.g007

Law, C.W., Chen, Y., Shi, W., Smyth, G.K., 2014. voom: Precision weights unlock linear model analysis tools for RNA-seq read counts. Genome Biol 15, R29. doi:10.1186/gb-2014-15-2-r29

Levin, D.E., 2011. Regulation of cell wall biogenesis in Saccharomyces cerevisiae: the cell wall integrity signaling pathway. Genetics 189, 1145–1175. doi:10.1534/genetics.111.128264

Longtine, M.S., McKenzie, A., Demarini, D.J., Shah, N.G., Wach, A., Brachat, A., Philippsen, P., Pringle, J.R., 1998. Additional modules for versatile and economical PCR-based gene deletion and modification in Saccharomyces cerevisiae. Yeast 14, 953–961. doi:10.1002/(SICI)1097-0061(199807)14:10<953::AID-YEA293>3.0.CO;2-U

Matsumoto, T.K., Ellsmore, A.J., Cessna, S.G., Low, P.S., Pardo, J.M., Bressan, R.A., Hasegawa, P.M., 2002. An osmotically induced cytosolic Ca2+ transient activates calcineurin signaling to mediate ion homeostasis and salt tolerance of Saccharomyces cerevisiae. J Biol Chem 277, 33075–33080. doi:10.1074/jbc.M205037200

Medina, D.L., Di Paola, S., Peluso, I., Armani, A., De Stefani, D., Venditti, R., Montefusco, S., Scotto-Rosato, A., Prezioso, C., Forrester, A., Settembre, C., Wang, W., Gao, Q., Xu, H., Sandri, M., Rizzuto, R., De Matteis, M.A., Ballabio, A., 2015. Lysosomal calcium signalling regulates autophagy through calcineurin and TFEB. Nat Cell Biol 17, 288–299. doi:10.1038/ncb3114

Minas, C., Waddell, S.J., Montana, G., 2011. Distance-based differential analysis of gene curves. Bioinformatics 27, 3135–3141. doi:10.1093/bioinformatics/btr528

Mizunuma, M., Hirata, D., Miyahara, K., Tsuchiya, E., Miyakawa, T., 1998. Role of calcineurin and Mpk1 in regulating the onset of mitosis in budding yeast. Nature 392, 303–306. doi:10.1038/32695

Mizunuma, M., Hirata, D., Miyaoka, R., Miyakawa, T., 2001. GSK-3 kinase Mck1 and calcineurin coordinately mediate Hsl1 down-regulation by Ca2+ in budding yeast. EMBO J 20, 1074–1085. doi:10.1093/emboj/20.5.1074

Morgan, D.O., 2007. The Cell Cycle. New Science Press.

Muzzey, D., Gómez-Uribe, C.A., Mettetal, J.T., van Oudenaarden, A., 2009. A systems-level analysis of perfect adaptation in yeast osmoregulation. Cell 138, 160–171. doi:10.1016/j.cell.2009.04.047

Ono, K., Han, J., 2000. The p38 signal transduction pathway: activation and function. Cell. Signal. 12, 1–13.

Orlando, D.A., Lin, C.Y., Bernard, A., Wang, J.Y., Socolar, J.E.S., Iversen, E.S., Hartemink, A.J., Haase, S.B., 2008. Global control of cell-cycle transcription by coupled CDK and network oscillators. Nature 453, 944–947. doi:10.1038/nature06955

Piao, H., MacLean Freed, J., Mayinger, P., 2012. Metabolic activation of the HOG MAP kinase pathway by Snf1/AMPK regulates lipid signaling at the Golgi. Traffic 13, 1522–1531. doi:10.1111/j.1600-0854.2012.01406.x

Pic-Taylor, A., Darieva, Z., Morgan, B.A., Sharrocks, A.D., 2004. Regulation of Cell Cycle-Specific Gene Expression through Cyclin-Dependent Kinase-Mediated Phosphorylation of the Forkhead Transcription Factor Fkh2p. Mol Cell Biol 24, 10036–10046. doi:10.1128/MCB.24.22.10036-10046.2004

Pramila, T., Wu, W., Miles, S., Noble, W.S., Breeden, L.L., 2006. The Forkhead transcription factor Hcm1 regulates chromosome segregation genes and fills the S-phase gap in the transcriptional circuitry of the cell cycle. Genes & Development 20, 2266–2278. doi:10.1101/gad.1450606

Reynolds, D., 2003. Recruitment of Thr 319-phosphorylated Ndd1p to the FHA domain of Fkh2p requires Clbkinase activity: a mechanism for CLB cluster gene activation. Genes & Development 17, 1789–1802. doi:10.1101/gad.1074103

Rothstein, R., 1991. Targeting, disruption, replacement, and allele rescue: integrative DNA transformation in yeast. Meth Enzymol 194, 281–301.

Sadasivam, S., DeCaprio, J.A., 2013. The DREAM complex: master coordinator of cell cycle-dependent gene expression. Nat Rev Cancer 13, 585–595. doi:10.1038/nrc3556

Schmitt, M.E., Brown, T.A., Trumpower, B.L., 1990. A rapid and simple method for preparation of RNA from Saccharomyces cerevisiae. Nucleic Acids Res 18, 3091–3092.

Sotelo, J., Rodríguez-Gabriel, M.A., 2006. Mitogen-activated protein kinase Hog1 is essential for the response to arsenite in Saccharomyces cerevisiae. Eukaryotic Cell 5, 1826–1830. doi:10.1128/EC.00225-06

Spellman, P.T., Sherlock, G., Zhang, M.Q., Iyer, V.R., Anders, K., Eisen, M.B., Brown, P.O., Botstein, D., Futcher, B., 1998. Comprehensive identification of cell cycle-regulated genes of the yeast Saccharomyces cerevisiae by microarray hybridization. Mol Biol Cell 9, 3273–3297.

Thorsen, M., Di, Y., Tängemo, C., Morillas, M., Ahmadpour, D., Van der Does, C., Wagner, A., Johansson, E., Boman, J., Posas, F., Wysocki, R., Tamás, M.J., 2006. The MAPK Hog1p modulates Fps1p-dependent arsenite uptake and tolerance in yeast. Mol Biol Cell 17, 4400–4410. doi:10.1091/mbc.e06-04-0315

Vaeth, M., Feske, S., 2018. NFAT control of immune function: New Frontiers for an Abiding Trooper. F1000Res 7, 260–13. doi:10.12688/f1000research.13426.1

Willis, N., Rhind, N., 2009. Mus81, Rhp51(Rad51), and Rqh1 Form an Epistatic Pathway Required for the S-Phase DNA Damage Checkpoint. Mol Biol Cell 20, 819–833. doi:10.1091/mbc.E08-08-0798

Winkler, A., Arkind, C., Mattison, C.P., Burkholder, A., Knoche, K., Ota, I., 2002. Heat stress activates the yeast high-osmolarity glycerol mitogen-activated protein kinase pathway, and protein tyrosine phosphatases are essential under heat stress. Eukaryotic Cell 1, 163–173. doi:10.1128/EC.1.2.163-173.2002

Yokoyama, H., Mizunuma, M., Okamoto, M., Yamamoto, J., Hirata, D., Miyakawa, T., 2006. Involvement of calcineurin-dependent degradation of Yap1p in Ca2+-induced G2 cell-cycle regulation in Saccharomyces cerevisiae. EMBO Rep. doi:10.1038/sj.embor.7400647

Yoshimoto, H., Saltsman, K., Gasch, A.P., Li, H.X., Ogawa, N., Botstein, D., Brown, P.O., Cyert, M.S., 2002. Genome-wide analysis of gene expression regulated by the calcineurin/Crz1p signaling pathway in Saccharomyces cerevisiae. J Biol Chem 277, 31079–31088. doi:10.1074/jbc.M202718200

Zhang, X., Cheng, X., Yu, L., Yang, J., Calvo, R., Patnaik, S., Hu, X., Gao, Q., Yang, M., Lawas, M., Delling, M., Marugan, J., Ferrer, M., Xu, H., 2016. MCOLN1 is a ROS sensor in lysosomes that regulates autophagy. Nature Communications 7, 12109. doi:10.1038/ncomms12109

