## Supplemental Information for "The coordinate actions of calcineurin and Hog1 mediate the response to cellular stress through multiple nodes of the cell cycle network"

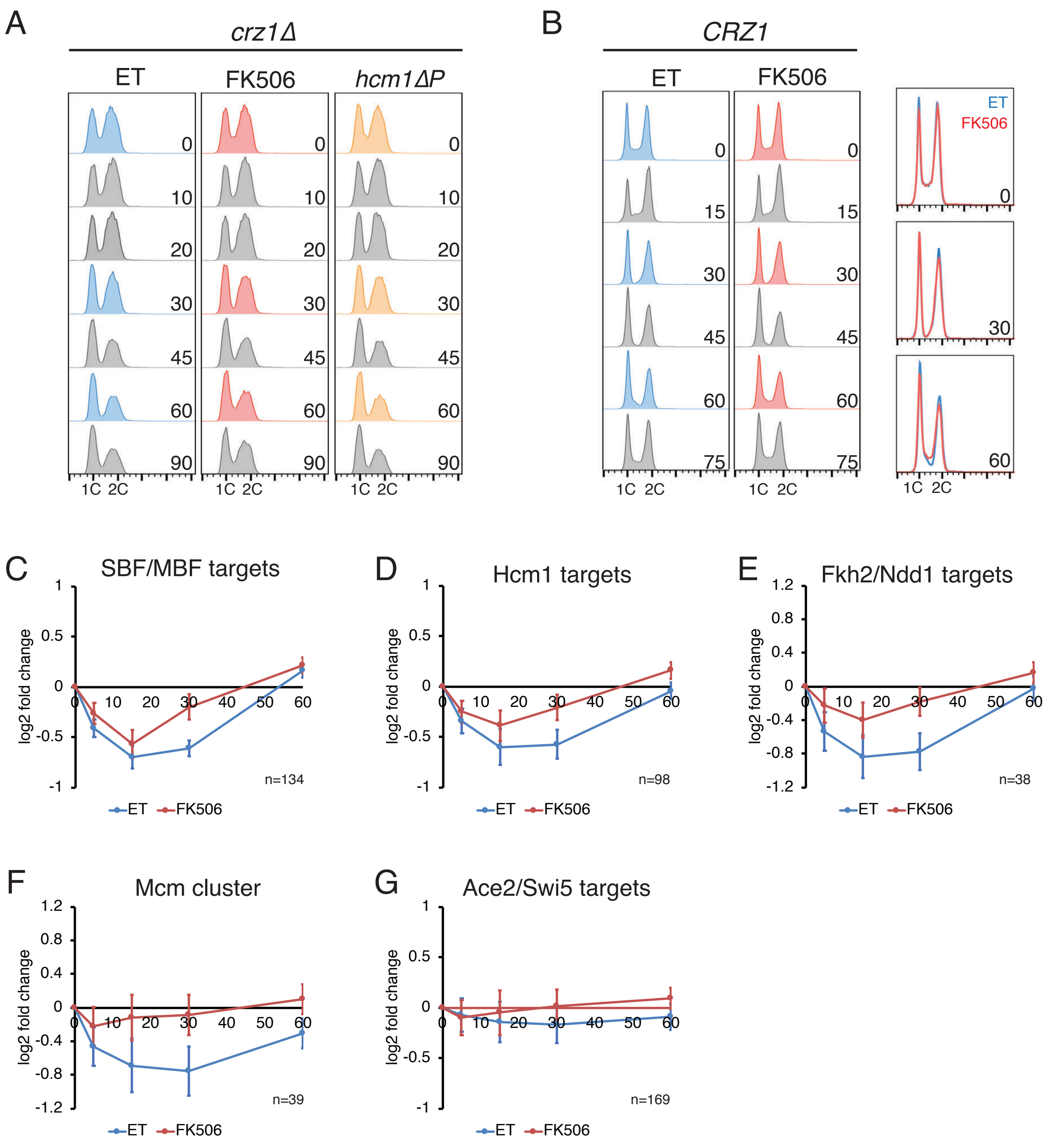


**Figure S1. Regulation of the cell cycle by CN. (A)** FACS plots from representative $CaCl_2$ time course in *crz1Δ* cells, in support of Figure 1A. **(B)** $CaCl_2$ time course in wild type (*CRZ1* proficient) cells. Wild-type cells were treated with ET or FK506 for 15 minutes before the addition of 200mM $CaCl_2$. Shown is DNA content as measured by flow cytometry at the indicated time points after $CaCl_2$ addition. Colored time points are overlaid to compare ET and FK506 samples in Figure 1A, left. **(C-G)** Average expression of the indicated groups of cell cycle-regulated genes after the addition of 200mM $CaCl_2$ to wild-type cells. Data is from (Yoshimoto et al., 2002). Number of genes in each cluster is indicated.

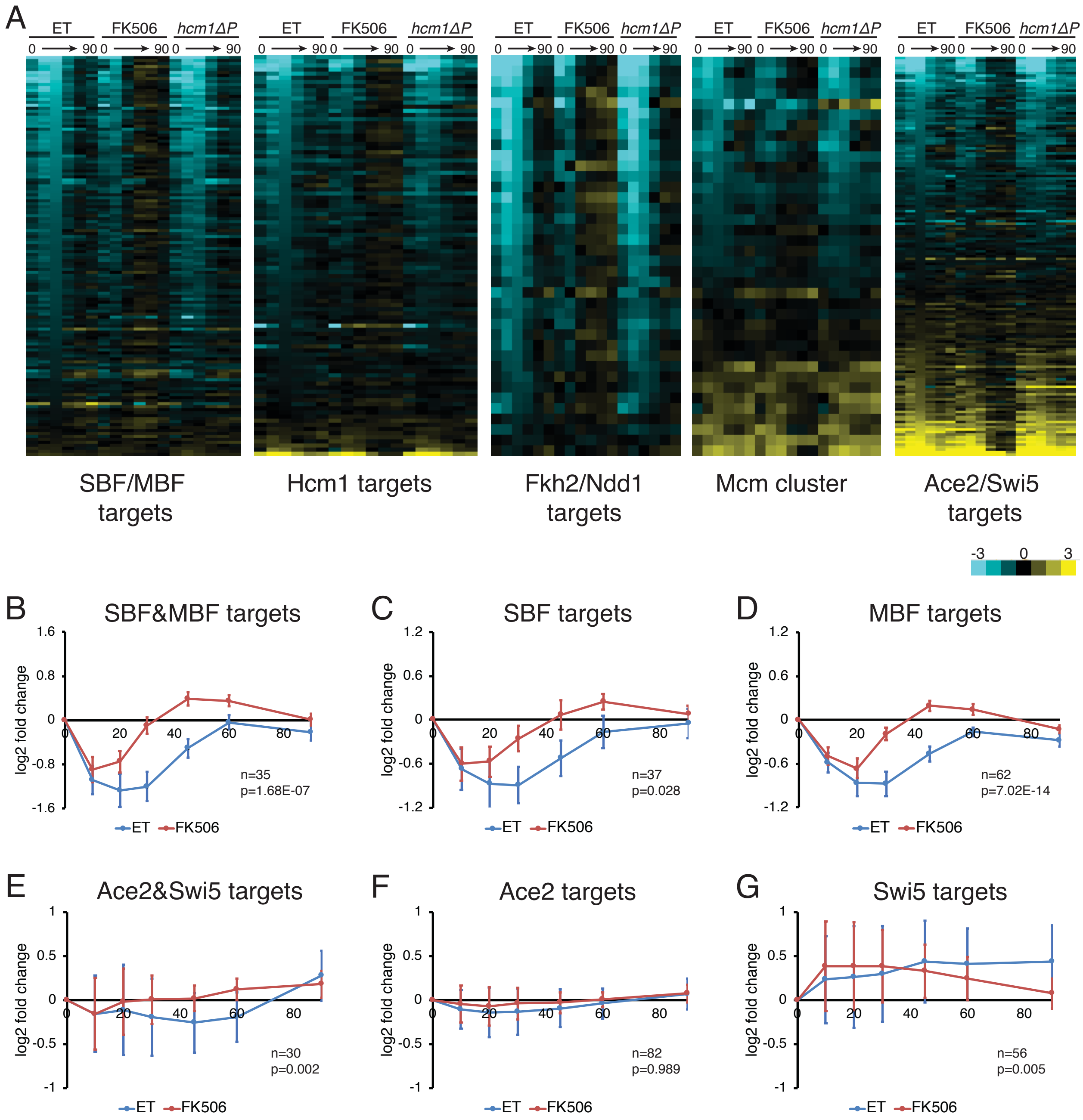


**Figure S2. Regulation of cell cycle gene expression by CN. (A)** Heat maps showing expression of all genes in each cluster that are averaged in Figure 1B-E. **(B-D)** Average expression of SBF/MBF target genes from Figure 1B, divided into subsets of genes regulated by both SBF and MBF (B), only SBF (C), or only MBF (D). Error bars indicate 95% confidence intervals. **(E-G)** Average expression of Ace2/Swi5 target genes from Figure 1F, divided into subsets of genes that are regulated by both Ace2 and Swi5 (E), only Ace2 (F), or only Swi5 (G). Error bars represent 95% confidence intervals. Number of genes (n) and the adjusted p-value indicating the significance of the difference between ET and FK506 curves are indicated.

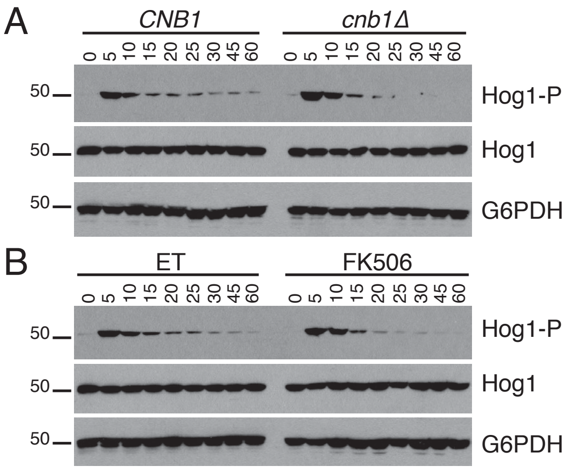


**Figure S3. CN maintains Hog1 activation in wild-type cells. (A)** Hog1 activation in wild-type cells is regulated by CN. Wild-type (*CRZ1*) cells or *cnb1Δ* cells were treated with 200mM $CaCl_2$ for the indicated number of minutes and Hog1 activation monitored by Western blot (Hog1-P). Total Hog1 and G6PDH (loading control) are shown. **(B)** Same as in (A) except wild type (*CRZ1 CNB1*) cells were pre-treated with ET buffer or FK506 for 15 minutes before the addition of $CaCl_2$.

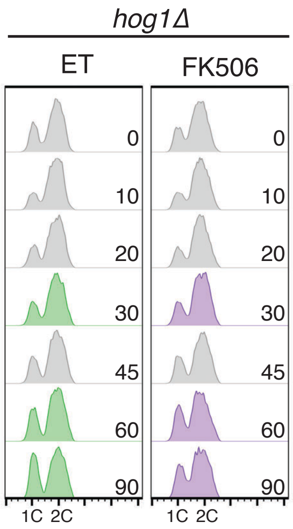


**Figure S4. Hog1 is required for the initial cell cycle arrest in response to $CaCl_2$ stress.** FACS plots from representative $CaCl_2$ time course in *crz1Δ hog1Δ* cells, in support of Figure 3A. Colored time points are overlaid in Figure 3A.

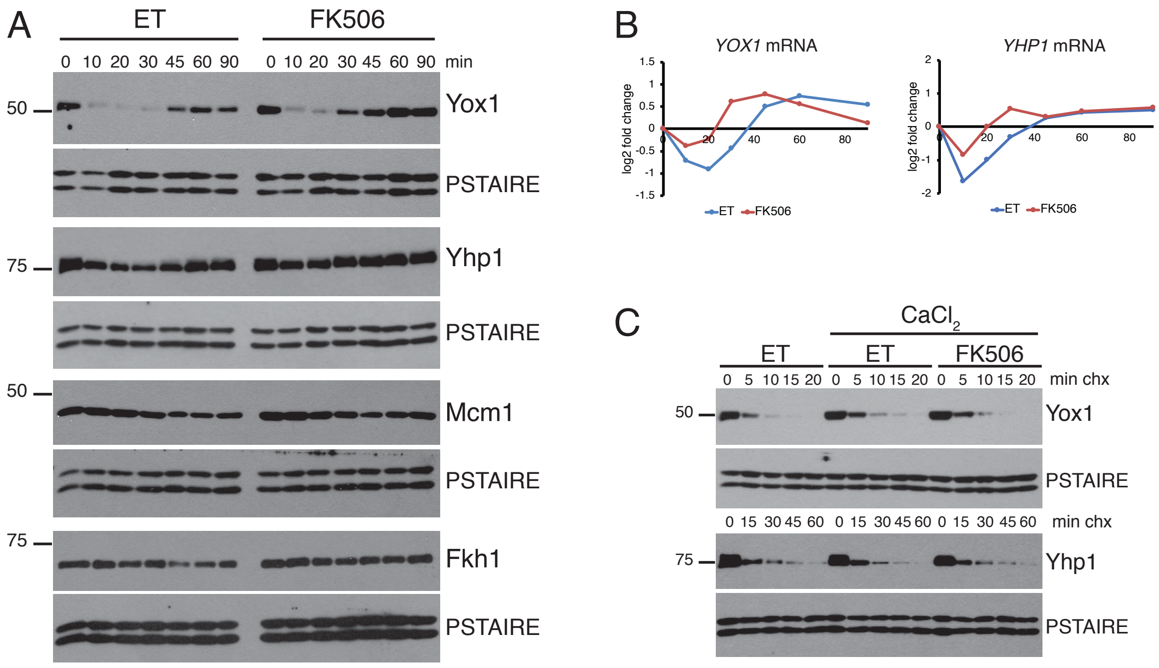


**Figure S5. Regulation of additional G2/M TFs by CN. (A)** Expression of TF proteins in response to $CaCl_2$. Strains expressing the indicated tagged TFs were pretreated with ET buffer or FK506 for 15 minutes before the addition of 200mM $CaCl_2$. Samples were collected for Western blotting at the indicated time points. Western blots were performed for a 3V5 tag on Fkh1, Mcm1, and Yox1 or a 13MYC tag on Yhp1. For all experiments PSTAIRE blots are shown as a loading control. **(B)** Expression of TF mRNAs in response to $CaCl_2$. Shown are log2 fold change values, compared to the 0-minute time point, from RNA-seq experiments described in Figure 1. **(C)** Cycloheximide-chase assays of the indicated TF proteins. Cells expressing tagged TF proteins from (A) were pretreated with ET buffer or FK506 for 10 minutes, 200mM $CaCl_2$ was added for an additional 5 minutes, then cycloheximide was added (0 minutes) and samples collected at the indicated time points for Western blot.

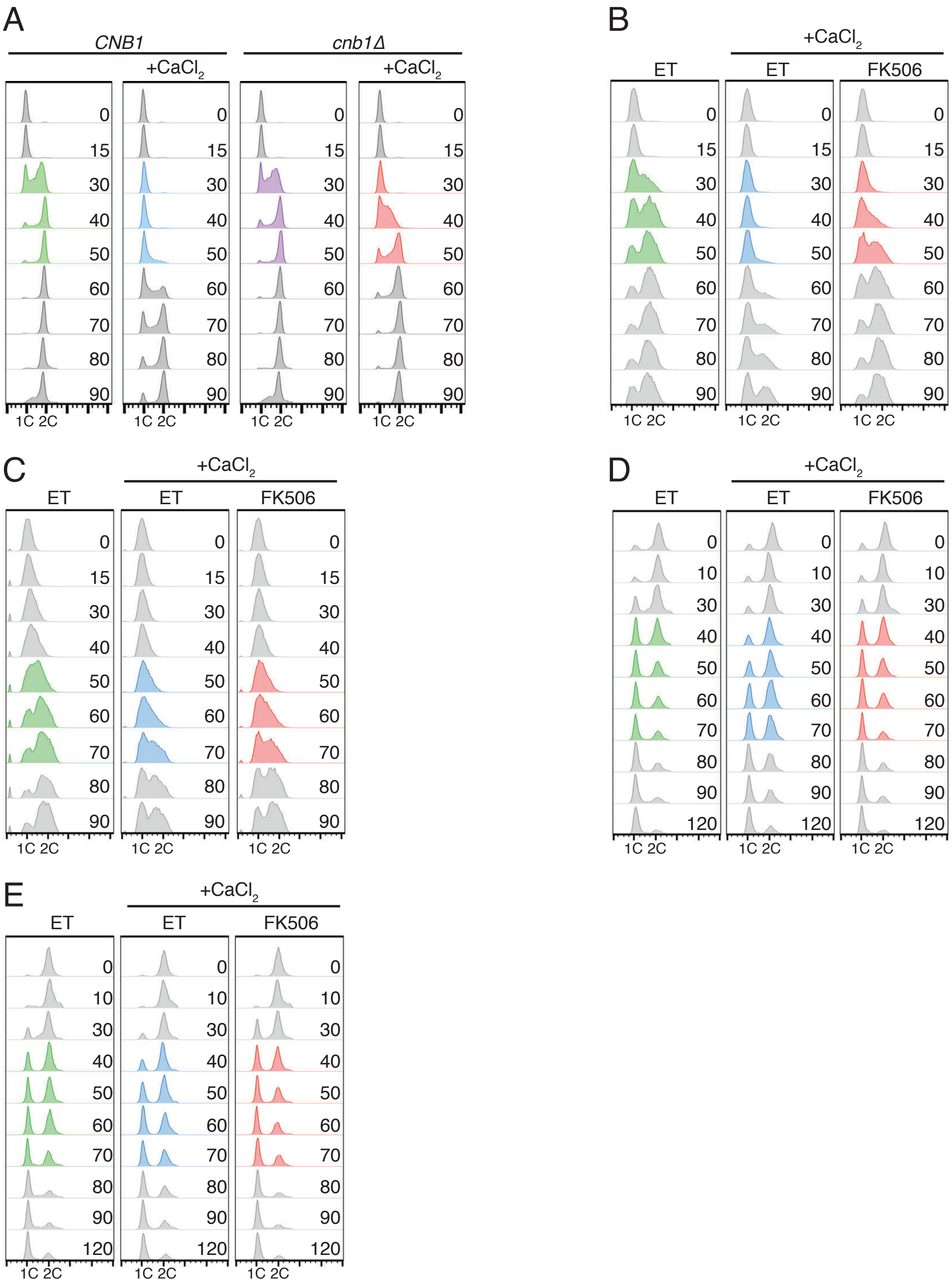


**Figure S6. Representative FACS plots to support Figure 6.** Representative FACS data from experiments that are quantified in Figure 6. In each panel time points that display the largest differences are highlighted in color. **(A)** Data from Figure 6B. **(B)** Data from Figure 6C. **(C)** Data from Figure 6D. **(E)** Data from Figure 6E. **(F)** Data from Figure 6G.

**Table S1. CN-regulated genes after 10 minutes $CaCl_2$ treatment**

| gene | ET/FK506 t=10* | BH adj p-value | gene cluster |
|---|---|---|---|
| *BAG7* | -1.856960889 | 0.000967319 | |
| *HAP4* | -1.163055122 | 8.23E-07 | |
| *REG2* | -1.150626613 | 0.042108723 | Ace2/Swi5 |
| *NRM1* | -1.141897458 | 0.007858729 | SBF/MBF |
| *ZEO1* | -1.0536199 | 0.029144541 | |
| *CLB2* | -1.031467576 | 0.000206166 | Fkh2/Ndd1 |
| *SUR7* | -0.949017459 | 0.000268532 | |
| *POG1* | -0.914706391 | 0.041214538 | |
| *CDC5* | -0.888550191 | 0.00199479 | Fkh2/Ndd1 |
| *FRK1* | -0.843561961 | 0.000975715 | Fkh2/Ndd1 |
| *COS111* | -0.834665457 | 0.012939887 | |
| *ALK1* | -0.796044016 | 0.00199479 | Fkh2/Ndd1 |
| *YNL058C* | -0.765573748 | 0.037184485 | Fkh2/Ndd1 |
| *SWI5* | -0.749489208 | 0.00199479 | Fkh2/Ndd1 |
| *TPO3* | -0.721860491 | 0.042108723 | Fkh2/Ndd1 |
| *MIG1* | -0.703374214 | 0.029144541 | |
| *IRC8* | -0.682302946 | 0.009439705 | Fkh2/Ndd1 |
| *PMA1* | -0.648080526 | 0.023655503 | |
| *SRC1* | -0.638086785 | 0.00199479 | Fkh2/Ndd1 |
| *ASE1* | -0.617232544 | 0.0027527 | Hcm1 |
| *YDR133C* | -0.6099885 | 0.048012708 | |
| *FKH2* | -0.609401994 | 0.029144541 | Hcm1 |
| *ACE2* | -0.605854216 | 0.0333304 | Fkh2/Ndd1 |
| *MRH1* | -0.56727327 | 0.042108723 | MCM cluster |
| *PHO90* | -0.553456027 | 0.019177709 | |
| *ECM33* | -0.469376696 | 0.029144541 | Hcm1 |
| *TSC10* | -0.421085289 | 0.041068935 | |
| *HST3* | -0.339988525 | 0.016264338 | Fkh2/Ndd1 |
| *MET22* | 0.481421259 | 0.028290041 | |
| *YAP5* | 0.651389782 | 0.04214145 | |
| *MET16* | 0.911730288 | 0.004058974 | |
| *ATG41* | 1.132156074 | 0.009224367 | |
| *STR3* | 1.572852266 | 0.00199479 | |

*Shown are log2 fold change values of ET-10min/FK506-10min

### Table S2. Strain table

| strain | genotype | figure |
|---|---|---|
| YHA146 | *MATa ura3Δ0 leu2Δ0 his3Δ0 met15Δ0 crz1Δ::KanMx HCM1-3HA-HIS3* | 1, 2C-D, 3B, 3E, 4B, 6C-E, S1A, S2, S5B, S6B-D |
| YHA147 | *MATa ura3Δ0 leu2Δ0 his3Δ0 met15Δ0 crz1Δ::KanMx hcm1-ΔPSIEIQ-3HA-HIS3* | 1, S1A, S2A |
| YCL34 | *MATa his3Δ ura3Δ leu2Δ met15Δ lys2Δ crz1Δ::LEU2 MCM1-3V5-KanMX* | 2A, S5A |
| YMF1 | *MATa his3Δ0 ura3Δ0 leu2Δ0 lys2Δ0 met15Δ0 crz1Δ::LEU2 cnb1Δ::KanMX HCM1-3HA-HIS3* | 2B, 6B, S6A |
| YMF7 | *MATa his3Δ0 ura3Δ0 leu2Δ0 crz1Δ::LEU2* | 2B, 6B, S6A |
| YCL37 | *MATa his3Δ ura3Δ leu2Δ met15Δ crz1Δ::LEU2 NDD1-3V5-KanMX* | 2E, 4A, 4C-D, 5B-D |
| YJB622 | *MATa ura3Δ0 leu2Δ0 his3Δ0 met15Δ0 crz1Δ::KanMx HCM1-3HA-HIS3 hog1Δ::LEU2* | 3A, 3C-D, 3F-G, S4 |
| YCL99 | *MATa his3Δ ura3Δ leu2Δ met15Δ crz1Δ::LEU2 FKH2-3FLAG-Hyg* | 4A, 4C-D, 5B-C |
| YMF42 | *MATa his3Δ ura3Δ leu2Δ met15Δ crz1Δ::LEU2 swe1Δ::URA3 NDD1-3V5-KanMX* | 5B-D, 6F, S6E |
| YMF44 | *MATa his3Δ ura3Δ leu2Δ met15Δ crz1Δ::LEU2 hog1Δ::HIS3 NDD1-3V5-KanMX* | 5B-D |
| YMF45 | *MATa his3Δ ura3Δ leu2Δ met15Δ crz1Δ::LEU2 hog1Δ::HIS3 swe1Δ::URA3 NDD1-3V5-KanMX* | 5B-D |
| YMF41 | *MATa his3Δ ura3Δ leu2Δ met15Δ FKH2-3FLAG-Hyg crz1Δ::LEU2 swe1Δ::URA3* | 5B-C |
| YMF46 | *MATa his3Δ ura3Δ leu2Δ met15Δ crz1Δ::LEU2 hog1Δ::HIS3 FKH2-3LAG-Hyg* | 5B-C |
| YMF47 | *MATa his3Δ ura3Δ leu2Δ met15Δ crz1Δ::LEU2 hog1Δ::HIS3 swe1Δ::URA3 FKH2-3FLAG-Hyg* | 5B-C |
| YBL176 | *MATa ura3Δ0 leu2Δ0 his3Δ0 met15Δ0 HCM1-3HA-HIS3* | S1B |
| YMF15 | *MATa his3Δ0 ura3Δ0 met15Δ0 cnb1Δ::KanMX* | S3A |
| MW836a | *MATa ura3Δ0 leu2Δ0 his3Δ0 met15Δ0* | S3A-B |
| YCL31 | *MATa his3Δ ura3Δ leu2Δ met15Δ crz1Δ::LEU2 FKH1-3V5-KanMX* | S5A |
| YCL43 | *MATa his3Δ ura3Δ leu2Δ met15Δ crz1Δ::LEU2 YOX1-3V5-KanMX* | S5A, S5C |
| YCL53 | *MATa his3Δ ura3Δ leu2Δ met15Δ crz1Δ::LEU2 YHP1-13MYC-KanMx* | S5A, S5C |

All strains are in the BY4741 background.

### Table S3. Primer table

| gene | primer name | sequence |
|---|---|---|
| *ACT1* | ACT1fwd | ATGAAGTGTGATGTCGATGTCC |
| | ACT1rev | CCAATCCAGACGGAGTACTTTC |
| *STL1* | STL1fwd | TTGCGGTATTTCATCACTATCG |
| | STL1rev | CACTACAGTTGCGTGTCTGTCA |
| *YOX1* | YOX1fwd | ATTTGCTTTCATCACACACTCG |
| | YOX1rev | AACTCAATTCGTTTCTCCTTCG |
| *CLN1* | CLN1fwd | CTTTGGTTAGCGGCCAAAAC |
| | CLN1rev | AGAAAGGCGTGGAATACGAG |
| *CLB2* | CLB2fwd | TGCATGTACGGAAGATGAAATC |
| | CLB2rev | AAGAATTTGGCAAGAGTTCGAG |
| *CDC5* | CDC5fwd | ATGTCCCATCCAAATATCGTTC |
| | CDC5rev | AATTCCATTAATGAACCGTTGG |
| *CDC20* | CDC20fwd | GCGGTAACCGTTCTGTACTTTC |
| | CDC20rev | GGGACGTTTGGAGTTTCTAATG |
